# Imperfect innate immune antagonism renders SARS-CoV-2 vulnerable towards IFN-γ and -λ

**DOI:** 10.1101/2020.10.15.340612

**Authors:** Manuel Hayn, Maximilian Hirschenberger, Lennart Koepke, Jan H Straub, Rayhane Nchioua, Maria H Christensen, Susanne Klute, Caterina Prelli Bozzo, Wasim Aftab, Fabian Zech, Carina Conzelmann, Janis A Müller, Smitha Srinivasachar Badarinarayan, Christina M Stürzel, Ignasi Forne, Steffen Stenger, Karl-Klaus Conzelmann, Jan Münch, Daniel Sauter, Florian I Schmidt, Axel Imhof, Frank Kirchhoff, Konstantin MJ Sparrer

**Affiliations:** Institute of Molecular Virology, Ulm University Medical Center, 89081 Ulm, Germany; Gene Center & Max von Pettenkofer-Institute of Virology, Ludwig-Maximilians-Universität München, 81377 Munich, Germany; Biomedical Center, Zentrallabor für Proteinanalytik (Protein Analysis Unit), Department of Molecular Biology, Ludwig-Maximilians-Universität München, 82152 Planegg-Martinsried, Germany; Graduate School for Quantitative Biosciences (QBM), Ludwig-Maximilians-University of Munich, 81377 Munich, Germany; Institute for Medical Microbiology and Hygiene, Ulm University Medical Center, 89081 Ulm, Germany; Institute of Innate Immunity, University of Bonn, 53127 Bonn, Germany

**Author notes:** contributed equally, in alphabetical order (Manuel Hayn, Maximilian Hirschenberger, Lennart Koepke).

## Abstract

The innate immune system constitutes a powerful barrier against viral infections. However, it may fail because successful emerging pathogens, like SARS-CoV-2, evolved strategies to counteract it. Here, we systematically assessed the impact of 29 SARS-CoV-2 proteins on viral sensing, type I, II and III interferon (IFN) signaling, autophagy and inflammasome formation. Mechanistic analyses show that autophagy and type I IFN responses are effectively counteracted at different levels. For example, Nsp14 induces loss of the IFN receptor, whereas ORF3a disturbs autophagy at the Golgi/endosome interface. Comparative analyses revealed that antagonism of type I IFN and autophagy is largely conserved, except that SARS-CoV-1 Nsp15 is more potent in counteracting type I IFN than its SARS-CoV-2 ortholog. Altogether, however, SARS-CoV-2 counteracts type I IFN responses and autophagy much more efficiently than type II and III IFN signaling. Consequently, the virus is relatively resistant against exogenous IFN-α/β and autophagy modulation but remains highly vulnerable towards IFN-γ and -λ treatment. In combination, IFN-γ and -λ act synergistically, and drastically reduce SARS-CoV-2 replication at exceedingly low doses. Our results identify ineffective type I and II antagonism as weakness of SARS-CoV-2 that may allow to devise safe and effective anti-viral therapies based on targeted innate immune activation.

## INTRODUCTION

The severe acute respiratory syndrome coronavirus 2 (SARS-CoV-2) is a zoonotic, novel coronavirus that emerged at the end of 2019^1–3^. Infection with SARS-CoV-2 causes Coronavirus disease 2019 (COVID19)^4^. The virus rapidly spread all over the world owing to its higher transmission rates^5^ (R=2.5), as well as a lower morbidity and case fatality rates (CFR 3-4%)^6^ compared to previous epidemic coronaviruses like SARS-CoV-1 (R=2.0, CFR 11%) or MERS-CoV (R=0.9, CFR 35%)^7–9^. However, its pathogenicity is still much higher than that of ‘common cold’ CoVs such as HKU1 and 229E^10^ and to date SARS-CoV-2 has caused more than a millions deaths (https://coronavirus.jhu.edu/map.xhtml).

Upon infection of a target cell, CoVs are recognized by innate immune sensors, for example via RIG-like receptors (RLRs)^11^, which activate cell-intrinsic innate immune defenses (hereafter referred to as the innate immune system)^12,13^, such as interferon (IFN) responses, inflammasome activation and autophagy^14^. However, the exact ligand triggering the response is unknown. Activation of RLRs induces signaling cascades that ultimately lead to the release of IFNs and other pro-inflammatory cytokines as well as induction of anti-viral effectors^15^. Released cytokines are subsequently also recognized by neighboring cells and induce an antiviral transcriptional response. Thus, both the infected cell and noninfected neighboring cells are set in an anti-viral state^16,17^ eventually limiting viral spread. Other branches of the innate immune system, such as autophagy, are activated during CoV infections as well^18,19^. Autophagy is capable of targeting viral components or even whole viruses for lysosomal degradation^20,21^ and SARS-CoV-2 has evolved to block autophagic turnover^18^. Eventually activation of innate immunity recruits and stimulates the adaptive immune system ultimately facilitating elimination of the virus^22,23^. Notably, inborn defects in innate immunity or auto-antibodies against IFNs are associated with high frequencies of severe COVID19 cases, suggesting that innate defense mechanisms play a major role in immune control of SARS-CoV-2^24,25^. Notably, SARS-CoV-2 infections show higher numbers of subclinical, asymptomatic infections (up to 80%^6^) compared to previous epidemic CoVs such as SARS-CoV^10^. Indeed, recent evidence suggest that SARS-CoV-2 can be more efficiently antagonized by IFNs than SARS-CoV-1 *in vitro^26^.* However, the underlying reasons for differences in IFN susceptibility between SARS-CoV-2 and SARS-CoV-1 are currently not fully understood.

Recent reports demonstrated that infection with SARS-CoV-2 induces an imbalanced innate immune response, indicating manipulation by SARS-CoV-2^27,28^. Proteomics analysis of selected SARS-CoV-2 proteins revealed that innate immune activation is perturbed on multiple levels^27^. For example, it was suggested that ORF3a inhibits autophagic turnover, ORF8 alters Integrin-TGFβ-EGFR-RTK signalling^27^ and ORF3b antagonizes type I IFN induction by a yet unknown mechanism^29^. In addition, the SARS -CoV −2 non-structural protein 1 (Nsp1) shuts down cellular translation including the cytokine – mediated innate immune response^30^. Analysis of the interplay between SARS-CoV-2 proteins and IFN-β induction and signaling revealed that at least 8 SARS-CoV-2 proteins interfere with type I IFN signalling^31,32^. Among them is ORF6, which was suggested to interfere with nuclear trafficking of transcription factors thereby impairing gene induction^32,33^. However, so far only type I IFN signaling was analyzed in some detail and our knowledge how SARS-CoV-2 manipulates innate immunity is far from being complete.

Currently, treatment with IFNs is explored in clinical trials against SARS-CoV-2^34^. However, patients receiving immunomodulatory therapy with IFNs generally suffer from severe side-effects including psychological symptoms such as depression^35–37^. Novel strategies which activate the immune system but reduce inflammation and lower doses of cytokines are required^38^. Thus, analyzing how SARS-CoV-2 antagonizes innate immunity may give valuable clues on viral vulnerabilities that might be exploited for effective and safe therapeutic immune control.

Here, we systematically analyzed the impact of 29 SARS-CoV-2 encoded proteins^29,39,40^ on the major branches of the cell-intrinsic innate immune system: IFN induction, IFN/pro-inflammatory cytokine signaling and autophagy. This identified Nsp1, Nsp3, Nsp5, Nsp10, Nsp13, Nsp14, ORF3a, ORF6, ORF7a and ORF7b as the major innate immune antagonists encoded by SARS-CoV-2. Interference with innate immune activation is achieved by using a diverse, synergistic, set of mechanisms ranging from downregulation of IFN receptor expression by Nsp14 to blockage of autophagy via fragmentation of the trans-Golgi network by the viral proteins ORF3a and ORF7a. Strikingly, our data indicate that Nsp15 of both RaTG13 CoV and SARS-CoV-2 counteract type I IFN induction and signaling much less efficiently than SARS-CoV-1 Nsp15. Taken together our analyses revealed that IFN-γ and IFN-λ1 pathways are antagonized the least, and consequently treatment with these two cytokines is most potent against SARS-CoV-2. Combined IFN treatment at very low doses potentiates the individual anti-viral effect and can be further improved by anti-inflammatory autophagy activation. Thus, our results provide a plausible explanation why SARS-CoV-2 is more susceptible against IFN treatment than SARS-CoV-1 and indicate that combination of IFN-γ and IFN-λ1 is an effective anti-SARS-CoV-2 approach.

## RESULTS

### A variety of SARS-CoV-2 proteins antagonize innate immune pathways

To systematically examine how SARS-CoV-2 manipulates innate immunity, we used Strep II-tagged expression constructs coding for 28 of the 30 currently reported SARS-CoV-2 proteins (Nsp1, Nsp2, Nsp4, Nsp5, Nsp6, Nsp7, Nsp8, Nsp9, Nsp10, Nsp11, Nsp12, Nsp13, Nsp14, Nsp15, Nsp16, S, ORF3a, ORF3c, E, M, ORF6, ORF7a, ORF7b, ORF8, ORF9b, N, ORF9c and ORF10) (Fig. 1a). In addition, we examined untagged Nsp3. Expression of all proteins was confirmed by western blotting (Supplementary Fig. 1a) and immunofluorescence analyses (Supplementary Fig. 1b). The impact of all 29 viral proteins on three major branches of innate immunity: IFN/pro-inflammatory cytokine induction via Rig-like receptors (Fig. 1b, Supplementary Fig. 1c), signaling (Fig. 1c, Supplementary Fig. 1d) and autophagy (Fig. 1d, Supplementary Fig. 1e) was analyzed by quantitative reporter assays.

**Figure 1:**
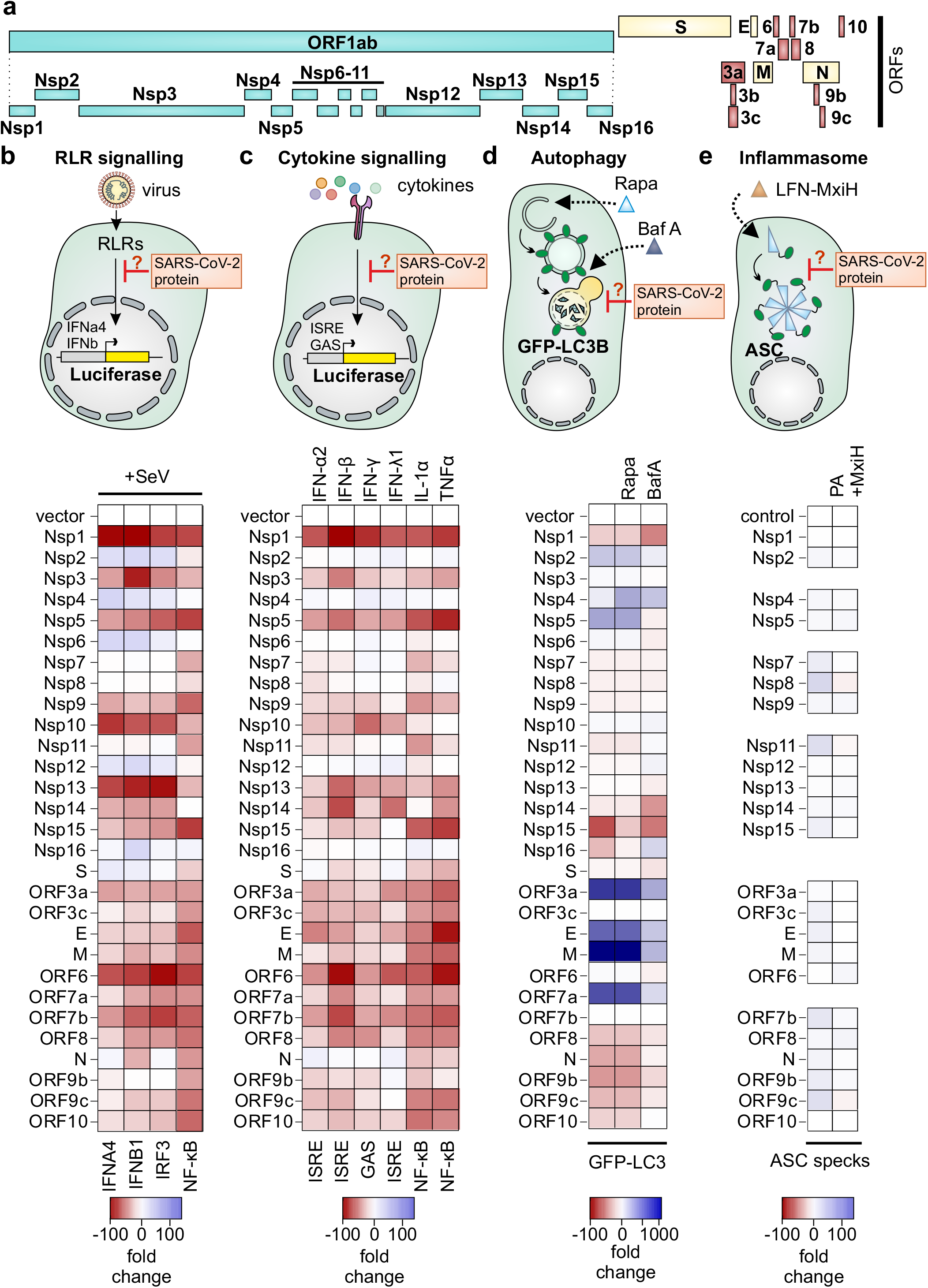
Systematic analysis of innate immune antagonism by SARS-CoV-2 proteins. **a**, Schematic depiction of the 30 SARS-CoV-2 encoded proteins in the order they appear in the genome. The polyprotein ORF1a(b) is (auto)proteolytically cleaved into 16 non-structural proteins (Nsp, turquoise). The structural proteins (yellow) are Spike (S), Membrane (M), envelope (E) and nucleoprotein (N). The set is complemented by the accessory proteins (red) ORF 3a, 3b, 3c, 6, 7a, 7b, 8, 9b, 9c and 10. **b-e**, Schematic depiction of the assay setup (top panel) and heatmap (red = inhibition, blue = induction) depicting modulation of innate immune pathways by overexpression of indicated SARS-CoV-2 proteins. Stimuli of the immune pathways are indicated. (a, b) Readout by Luciferase reporter gene assay (color represents the mean of n=3) using indicated promotor constructs in HEK293T cells, (c) autophagosome measurement by quantification of membrane-associated GFP-LC3B in HEK293T-GFP-LC3B cells (color represents the mean of n=4) or (d) Quantification of ASC specks in THP-1 cell lines by flow cytometry; cell lines doxycycline-inducible expressing the indicated transgenes were treated with Bacillus anthracis PA and LFn-MxiH to stimulate NLRC4 inflammasomes (color represents the mean of n=2). The vector/control is set to 1 (white). SeV, Sendai Virus. Rapa, Rapamycin. BafA, Bafilomycin A1, PA, protective antigen of *B. anthracis*, MxiH, Needle protein of *S. flexneri.*

Induction of type I IFNs (IFN-α and IFN-β) was monitored using a Firefly luciferase reporter controlled by the full IFNα4 promotor, the full IFN-β promotor, or isolated binding sites for the transcription factors IRF3 or NF-κB (Fig. 1b). All assays were normalized for cell viability (Supplementary Fig. 1f). HEK293T cells were infected with Sendai Virus, mimicking RLR activation by SARS-CoV-2. Nsp2, Nsp6 and Nsp12 slightly enhanced both IFN-α4 and IFN-β promotor induction as well as IRF3-dependent transcription (Fig. 1b). However, our analyses revealed that Nsp1, Nsp3, Nsp5, Nsp10, Nsp13, ORF6 and ORF7b are the major SARS-CoV-2 encoded antagonists of type I IFN induction (Fig. 1b).

Type I and III IFNs, such as IFN-α, IFN-β and IFN-λ1 culminates in the induction of genes with IFN response element (ISRE)-containing promotors^16^. Type II IFN-γ cause gene activation of gamma activated sequence (GAS) containing promotors. Pro-inflammatory cytokine signaling (TNFa and IL-1a) induce genes containing NF-ĸB sites in the promotor. Signaling of type I IFNs (IFN-α and IFN-β), Type II IFN (IFN-γ), type III IFN (IFN-λ1) and pro-inflammatory cytokine signaling (TNFa and IL-1a) was quantified using quantitative firefly luciferase reporters controlled by the respective promotors (Fig. 1c). Stimulation with IFN-α2 and IFN-β (Fig. 1c) revealed that activation of the ISRE promotor is strongly repressed by Nsp1, Nsp5, Nsp13, Nsp14, ORF6 and ORF7b. A similar set of viral proteins interfered with type II IFN-γ and type III IFN-λ1 signaling, albeit much weaker (mean inhibition 18% and 35%, respectively) compared to type I IFN signaling (mean inhibition 78% for IFN-α2 and 53% for IFN-β). Activation of NF-ĸB signaling by TNFa or IL-1α was potently inhibited by the SARS-CoV-2 Nsp1, Nsp5, Nsp15, ORF3a, E, M, ORF6 and ORF7b proteins. These analyses revealed that a similar set of proteins (Nsp1, Nsp5, Nsp15, ORF3a, E, M, ORF6 and ORF7b) antagonizes pro-inflammatory cytokine signaling.

Since induction of autophagy does not depend on *de novo* gene expression^41^, we monitored autophagy levels in SARS-CoV-2 protein expressing HEK293T cells by membrane-association of stably expressed GFP-LC3B, a hallmark of autophagy induction (Fig. 1d, Supplementary Fig. 1e)^42^. Autophagosome numbers under basal conditions were strongly increased in the presence of ORF3a, E, M and ORF7a suggesting either de novo induction of autophagy or blockage of turnover (Fig. 1d). Upon induction of autophagy using Rapamycin, a similar pattern was observed. To clarify whether these viral proteins induce autophagy or block turnover, leading to accumulation of GFP-LC3B positive vesicles, we treated cells with saturating amounts of Bafilomycin A1, which inhibits autophagic turnover. The increase of autophagosome numbers by ORF3a, E, M and ORF7a was drastically reduced compared to nonblocking conditions (Fig. 1d), indicating that these proteins block turnover, rather than induce it. Blockage of autophagy and co-expression of Nsp1 and Nsp14 induced cell death, which may be responsible for the low number of autophagosomes. Unexpectedly, in the presence of Nsp15 autophagosome numbers were consistently reduced, suggesting that it inhibits autophagy (Fig. 1d).

Inflammasome responses were analyzed in stable THP-1 cell lines expressing SARS-CoV-2 proteins upon doxycycline induction. To avoid any effects of transcription, assembly of ASC specks was quantified after cytosolic delivery of the NAIP/NLRC4 inflammasome activator *Shigella flexneri* needle protein MxiH using the anthrax toxin delivery system (Fig. 1 e)^43^. ASC speck assembly is typically followed by caspase-1 activation and release of pro-inflammatory IL-1β and IL-18^43,44^. Expression of the SARS-CoV-2 Nsp8, Nsp11 and ORF9c very weakly induced inflammasome activity in the absence of inflammasome activators, although counterselection against cells prone to aberrant inflammasome activation during selection cannot be ruled out. Activation of NLRC4 inflammasomes was not significantly antagonized by any viral protein.

Taken together, our analysis reveals that SARS-CoV-2 encodes multiple proteins that strongly antagonize innate immunity. Notably, there are differences in overall inhibition of the pathways with IFN-γ, IFN-λ1 as well as inflammasome activity signaling being only weakly antagonized. However, type-I IFN induction and signaling and autophagy are strongly repressed.

### SARS-CoV-2 proteins target autophagy and type-I IFN at multiple levels

To analyses mechanistically why type-I IFN and autophagy are potently counteracted by SARS-CoV-2, we aimed at identifying the steps that are targeted in these pathways. We focused on the top 5 inhibitors as identified in Fig. 1b-d. Nsp1 was removed from the analysis as it prevents translation in general^30^. To analyses IFN-β signaling, we monitored the levels of the type I IFN receptor, IFNAR using western blotting in HEK293T cells overexpressing Nsp5, Nsp13, Nsp14, ORF6 or ORF7b. Activation of the two major transcription factors of type I IFN signaling, STAT1 and STAT2 (Fig. 2a) was examined by phosphorylation status. Basal STAT1 and STAT2 levels were not significantly affected by all proteins tested (Fig. 2b, quantification in Supplementary Fig. 2a-c). (Fig. 2b). In the presence of Nsp5, activated STAT1 and to a lesser extend STAT2 accumulate (Fig. 2b and 2d, Supplementary Fig. 2a). ORF6 and ORF7b did not affect IFNAR levels or STAT1 expression or activation (Fig. 2b-d). This agrees with recent reports^26,45,46^ suggesting that ORF6 instead prevents trafficking of transcription factors. In the presence of Nsp14 and to a lesser extend for Nsp13 endogenous levels of IFNAR is prominently reduced (Fig. 2b, c). Consequently, phosphorylation of STAT1 was decreased upon Nsp14 co-expression (Fig. 2b, d).

**Figure 2:**
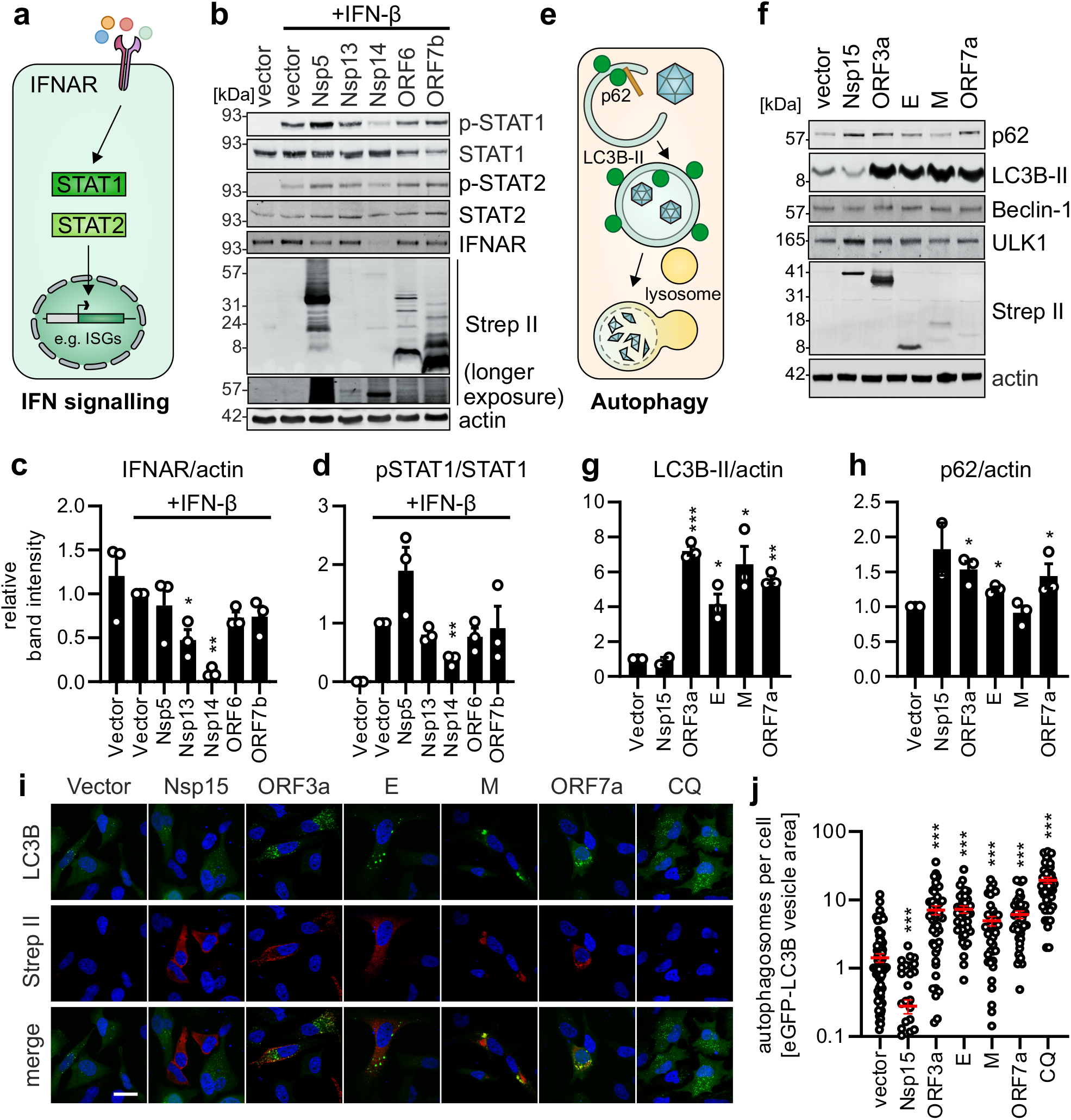
SARS-CoV-2 interferes with innate immunity at various levels. **a**, Schematic depiction of the type-I IFN signaling pathway. **b**, Exemplary immunoblot analysis showing activation of type-I IFN signaling markers using whole cell lysates (WCLs) of HEK293T cells expressing indicated proteins and stimulated with IFN-β (1000 U/mL, 45 min). Blots were stained with anti-pSTAT1, anti-STAT1, anti-pSTAT2, anti-STAT2, anti-IFNAR, anti-strep II and anti-actin. **c**, Quantification of the band intensities in (b) for IFNAR normalized to the band intensities of actin. Bars represent mean of n=3±SEM. **d**, Quantification of the band intensities in (b) for phospho-STAT1 (pSTAT1) normalized to the band intensities of actin. Bars represent mean of n=3±SEM. **e**, Schematic depiction of autophagy. **f**, Exemplary immunoblot analysis showing autophagy activity markers using WCLs of HEK293T cells expressing indicated proteins. Blots were stained with anti-SQSTM1/p62, anti-LC3B-II, anti-Beclin-1, anti-ULK1, anti-strep II and anti-actin. **g**, Quantification of the band intensities in (f) for LC3B-II normalized to the band intensities of actin. Bars represent mean of n=3±SEM. **h**, Quantification of the band intensities in (f) for p62 normalized to the band intensities of actin. Bars represent mean of n=3±SEM. **i**, Exemplary confocal laser scanning microscopy images of autophagy activation via GFP-LC3B (green) puncta formation. Indicated strep II-tagged SARS-CoV-2 proteins (red) were overexpressed in HeLa GFP-LC3B cells (green). CQ, Chloroquine (4 h 10 μM) was used as a positive control. Nuclei, DAPI (blue). Scale bar, 25 μM. **j**, Quantification by area of GFP-LC3B puncta divided by cell number from the images in (i). Bars represent the mean of n=38-100 cells±SEM.

Upon activation of autophagy, cytoplasmic MAP1LC3B (LC3B) is proteolytically processed and lipidated (LC3B-II) to decorate autophagosomal membranes^41,42^. Upon fusion of autophagosomes with lysosomes, the autophagic receptor p62 is degraded (autophagy turnover, Fig. 2e). We analyzed the effect of the top 5 autophagy modulating SARS-CoV-2 proteins: Nsp15, ORF3a, E, M and ORF7a (Fig. 1d) on autophagy markers. Levels of Beclin-1 and ULK1, which parts of the core machinery of autophagy initiation remained constant (Fig. 2f, Supplementary Fig. 2d and 2e). Overexpression of Nsp15 leads to a very slight decrease of LCB3-II but accumulation of p62, suggesting that Nsp15 blocks induction of autophagy (Fig. 2f and 2g-h). In line with this, the number of GFP-LC3B-puncta (=autophagosomes) per cell in HeLa-GFP-LC3B cells is reduced upon Nsp15 expression to almost 0 (Fig. 2i, j). In the presence of ORF3a, E and ORF7a, the levels of processed LC3B (LC3B-II) were 4-to 7-fold increased (Fig. 2g), and p62 levels are approximately 1.5-fold increased (Fig. 2h). This indicates that these three viral proteins block autophagic turnover. Consequently, the number of autophagosomes is 10-fold increased upon ORF3a, E, M or ORF7a expression (Fig. 2i, j). Curiously, while accumulation of LC3B-II indicates that M blocks autophagic turnover or induces autophagy, the levels of p62 are not significantly altered in the presence of M (Fig. 2f, h). Notably, overexpression of M resulted in an accumulation of LC3B in the perinuclear space, whereas for all other viral proteins autophagosomes are normally distributed (Fig. 2i, j).

Taken together, our data demonstrates that SARS-CoV-2 synergistically targets type-IFN signaling and autophagy. The major type I IFN antagonists Nsp5, Nsp13, Nsp14, ORF6 or ORF7b block the signaling cascade at different levels. E, ORF3a and ORF7a use similar mechanism to block autophagic turnover, while M may have evolved a different mechanism and Nsp15 inhibits de novo autophagy induction.

### ORF3a and ORF7a perturb the late-endosomal/trans-Golgi network

Our data showed that ORF3a and ORF7a are the most potent autophagy antagonists of SARS-CoV-2 (Fig. 1d, Fig. 2f-j). To determine their molecular mechanism(s), we performed proteome analysis of HEK293T cells overexpressing SARS-CoV-2 ORF3a and ORF7a (Supplementary Fig. 3a). As a control, we used S, Nsp1 and Nsp16 overexpressing cells which show little to no effect on autophagy (Fig. 1d). In addition, we analyzed the proteome of Caco-2 cells infected with SARS-CoV-2 for 24 or 48 h. Fold changes compared to vector transfected or non-infected controls were calculated (Fig. 3a, b, Supplementary Fig. 3b-e, Supplementary Table 1). Analysis of the data revealed that in the presence of Nsp1, cellular proteins with a short half-life are markedly reduced (Supplementary Fig. 3f)^47^. This supports our previous finding that Nsp1 globally blocks translation^30^ and confirming the validity of the proteome analysis. PANTHER-assisted Gene Ontology Analysis of the proteins regulated more than 4-fold by the overexpression of individual SARS-CoV-2 proteins revealed that ORF3a and ORF7a target the late endosome pathway (GO:0005770) (Fig. 3c, Supplementary Table 2). A similar analysis for the SARS-CoV-2 samples showed that the late endosome pathway is also affected during the genuine infection. Thus, we had a closer look at the subcellular localization of ORF3a and ORF7a and their effect on intracellular vesicles. In line with the proteome analysis, ORF7a and ORF3a both localized to the late endosomal compartment, co-localizing with the marker Rab9 (Fig. 3d,e). In contrast, localization to Rab5a-positive early endosomes was not apparent (Supplementary Fig. 3g). Disturbance of the integrity of the trans-Golgi network (TGN) at the interface with the late endosomes^48,49^ by viral proteins is a well-known strategy to block autophagy^50^. Immunofluorescence analysis revealed that the localization of ORF3a or ORF7a partially overlap with a TGN marker (R = 0.5, Fig. 3g) indicating close proximity. ORF6, which is known to localize to the Golgi apparatus^45^ was used a positive control (R=0.7). Nsp8, which displayed a cytoplasmic localization was used as a negative control (R=0.3). Importantly, analysis of free TGN-marker positive vesicles in SARS-CoV-2 ORF3a or ORF7a expressing cells revealed that both viral proteins cause significant fragmentation of the TGN (Fig. 3f, h).

**Figure 3:**
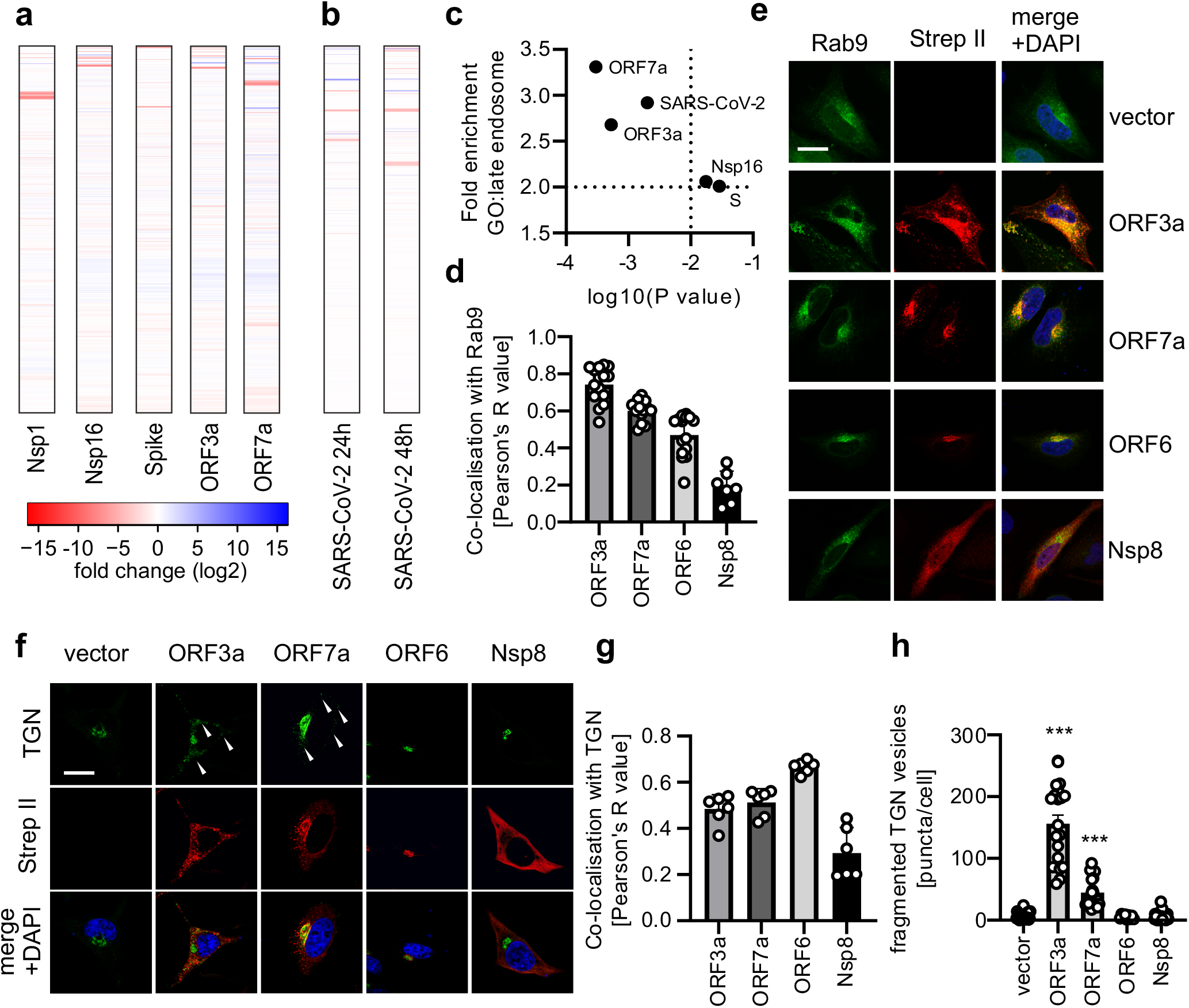
ORF3a and ORF7a disturb the trans-Golgi network/late endosome interface. **a**, Heatmap (red = downregulation, blue = upregulation) depicting the fold changes of cellular and viral proteins during overexpression of indicated single SARS-CoV-2 proteins in HEK293T cells or **b**, SARS-CoV-2 infection (MOI 1) of Caco-2 cells 24 or 48 h post infection as assessed by mass spectrometry. **c**, Scatter plots of log2 fold enrichment and p-value of the GO-Term ‘late endosome’ in protein sets regulated more than 4-fold upon expression of indicated viral protein (a) or SARS-CoV-2 infection (b). **d**, Quantification of co-localization by Pearson Correlation of Rab9 and indicated viral proteins in HeLa cells transiently transfected with the indicated viral protein and GFP-Rab9. Bars represent the mean of n=7-15 cells±SEM. **e**, Exemplary confocal microscopy images of HeLa cells transiently expressing indicated viral proteins (red) and a marker of late endosomes GFP-Rab9 (green). Cells were stained with anti-strep II (red). Nuclei, DAPI (blue). Scale bar, 10 μm. **f**, Exemplary confocal microscopy images of the quantification in (g) stained with anti-TGN46 (green) and anti-strep II (red). Nuclei, DAPI (blue). Scale bar, 10 μm. **g**, Pearson’s correlation indicating colocalization between TGN46 and the indicated viral proteins from the image in (f). Bars represent the mean of n=6 cells±SEM. h, Quantification of non-Golgi associated vesicles per cell as puncta/cell of (f). Bars represent the mean of n=15-25 cells ±SEM.

These data indicate, that both ORF3a and ORF7a disturb the proteome at the late endosomes eventually causing the TGN to fragment, which leads to a block of autophagic turnover^49–52^.

### SARS-CoV-2 Nsp15 is less potent in innate immune antagonism than SARS-CoV-1 Nsp15

To examine the conservation of innate immune antagonism, we functionally compared Nsp1, Nsp3, Nsp7, Nsp15, M, N, ORF3a, ORF6 and ORF7a of SARS-CoV-2, the closest related CoV, RaTG13-CoV and the previous highly pathogenic SARS-CoV-1. RaTG13-CoV was isolated from the intermediate host horseshoe bats *(Rhinolophus affinis).* The amino acid sequences of the different CoVs are largely conserved, with the exception of Nsp3, ORF3a and ORF6 (Fig. 4a) and are all expressed to similar levels as confirmed by western blotting (Supplementary Fig. 4a-i). Rabies virus P protein, Measles virus V protein and TRIM32 expression served as positive controls. Overall, proteins of SARS-CoV-1 and RaTG13 behave similar to their SARS-CoV-2 counterparts, suggesting that many functions are conserved. Importantly, however, this is not the case for Nsp15, Nsp3 and to a lesser extend ORF6 (Fig. 4a-c). SARS-CoV-1 ORF6 is about 4-fold less potent in antagonizing type I IFN signaling (Fig. 4b) but induces higher levels of autophagy (Fig. 4c). However, expression levels of SARS-CoV-1 ORF6 were also higher than that of its SARS-VoV-2 and RaTG13 counterparts (Supplementary Fig. 4g), which may explain the differences in activity. Differences between SARS-CoV, RaTG13 and SARS-CoV-2 Nsp3 were reanalyzed in a dose-dependent manner, and only in the range of 2-3-fold which may also explained by differential expression (Supplementary Fig. 4j).

**Figure 4:**
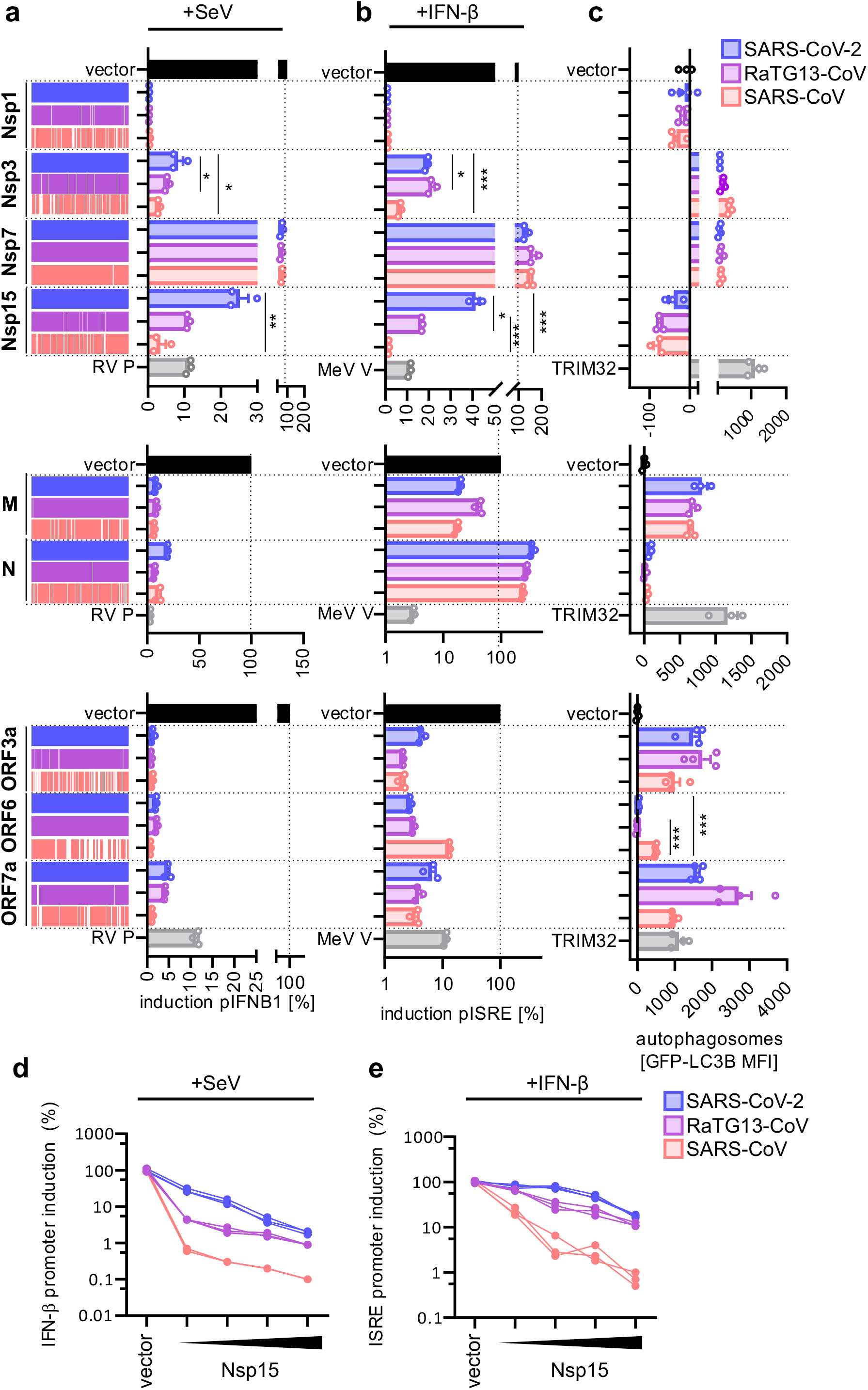
Conservation of innate immune antagonism between SARS-CoV-2, RaTG13-CoV and SARS-CoV. **a-c**, Immune activation of type-I IFN induction (a), type-I IFN signaling (b) or autophagy (c) in the presence of indicated proteins (Nsp1, Nsp3, Nsp7, Ndsp15, M, N, ORF3a, ORF6, ORF7a) of SARS-CoV-2 (blue), RaTG13-CoV (purple) or SARS-CoV-1 (red) assessed by IFN-β-promotor luciferase reporter gene assays stimulated with Sendai Virus (SeV, a). ISRE-promotor luciferase reporter gene assays stimulated with IFN-β (1000 U/ml, b). Membrane-associated GFP-LC3B (c) (n=4±SEM). Vector induction set to 100% (black). Controls, RABV P, MeV V or TRIM32 (grey). Bars represent the mean of n=3±SEM (a,b) or n=4±SEM (c). **d**, Dose dependent effect of SARS-CoV-2, RaTG13-CoV or SARS-CoV-1 Nsp15 expression on IFN-β induction stimulated with SeV (24 h). Quantification by IFN-β promotor dependent luciferase reporter activity. Lines represent one individual replicate. **e**, Dose dependent effect of Nsp15 expression on IFN-β signaling in HEK293T cells, stimulated with IFN-β (1000 U/ml, 8 h). Quantification by ISRE promotor dependent luciferase reporter activity. Lines represent one individual replicate.

The most striking, statistically significant difference was observed for Nsp15. SARS-CoV-1 Nsp15 is over 10-fold more potent in suppression of type I IFN induction and signaling than RaTG13 and SARS-CoV-1 Nsp15 (Fig. 4a, b). Notably, expression levels of SARS-CoV-2, RaTG13 and SARS-CoV-1 Nsp15 are similar, with SARS-CoV-1 Nsp15 even being slightly less expressed (Supplementary Fig. 4c). Notably, all Nsp15 variants still inhibit autophagy (Fig. 4c). Dose-dependent effect of SARS-CoV-2 Nsp15, RaTG13-CoV Nsp15 and SARS-CoV-1 Nsp15 on type I IFN induction (Fig. 4d) and signaling (Fig. 4e) showed that on average SARS-CoV2 Nsp15 performed 32-fold worse than SARS-CoV-1 Nsp15, and RaTG13 Nsp15 inhibited type I IFN induction 7.8-fold less (Fig. 4d). Similarly, SARS-CoV-1 Nsp15 outperformed RaTG13 and SARS-CoV-2 Nsp15 by 15-and 5.7-fold, respectively, in inhibition of type I IFN signaling (Fig. 4e).

Taken together, this data indicates, that while most IFN antagonist activities are conserved between SARS-CoV, RaTG13 and SARS-CoV-2, there is an exception: Nsp15 of SARS-CoV-1 is considerably more potent than SARS-CoV-2 Nsp15 in counteracting both IFN-β induction and signaling.

### Inefficient antagonism by SARS-CoV-2 proteins is predictive for efficient immune control

Our analyses revealed that several of the 29 SARS-CoV-2 proteins synergistically antagonize innate immune activation (Figs. 1–4), albeit with different efficiency. The mean inhibition of IFN-γ and IFN-X1 signaling was 18% and 35%, respectively, compared to type I IFN signaling with a mean inhibition of only 78% for IFN-α2 and 53% for IFN-β. Consequently, we assessed whether IFN-α2, IFN-β, IFN-γ and IFN-λ1 have a different impact on SARS-CoV-2 (Fig. 5a, Supplementary Fig. 5a, b). Treatment with the type I IFN-α2 was the least efficient. In contrast, at the same concentration IFN-γ (500 U/mL) reduced viral RNA in the supernatant almost 300-fold more efficiently. All agents caused little if any cytotoxic effects (Supplementary Fig. 5c). Altogether, we observed a good correlation (r= 0.89) between average inhibition of the respective signaling pathway (Fig. 1c) antagonized by the 29 SARS-CoV-2 proteins and IFN susceptibility at 5 U/mL (Fig. 5b).

**Figure 5:**
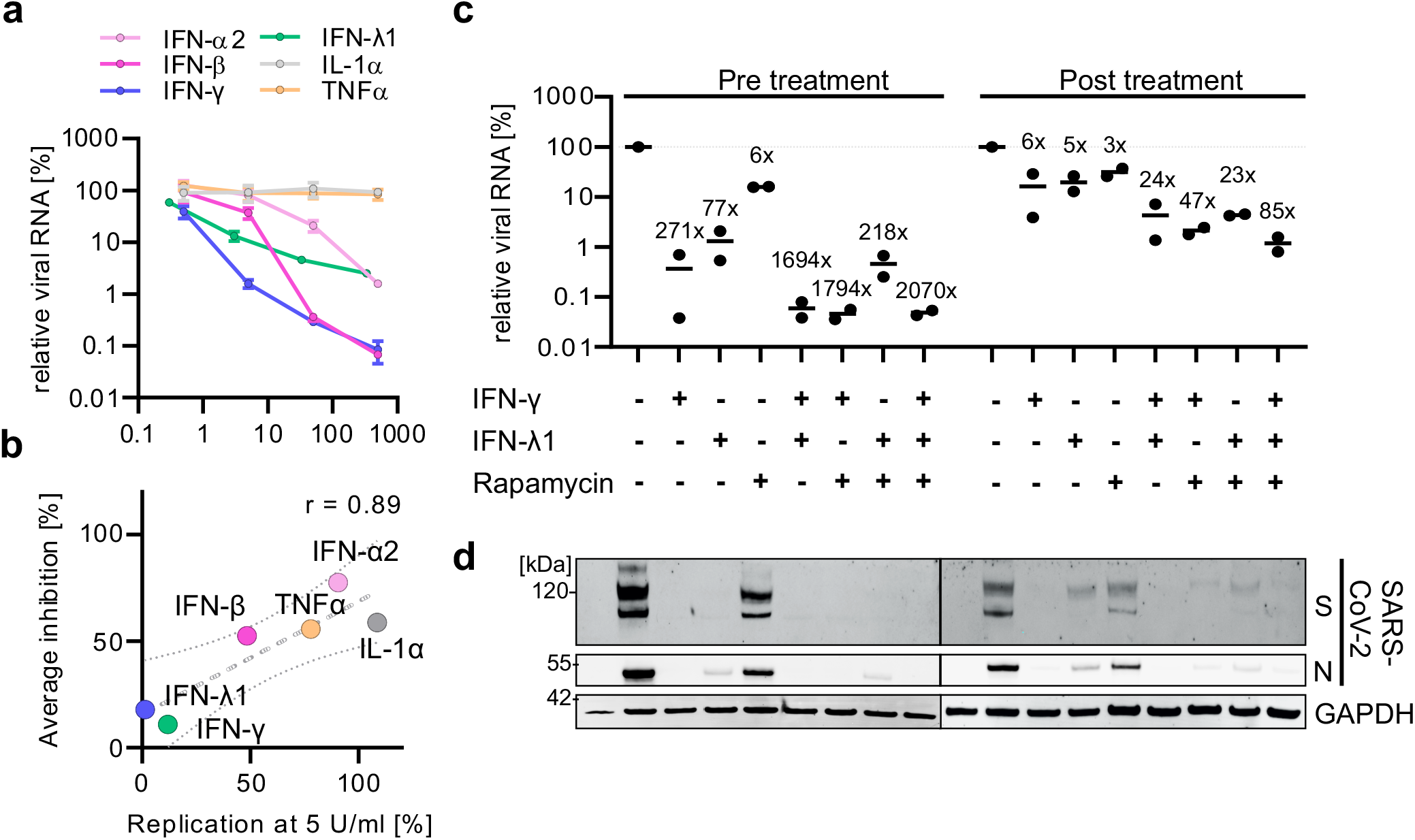
Innate immune activation as an anti-viral approach. **a**, SARS-CoV-2 N RNA in the supernatant of SARS-CoV-2 (MOI 0.05, 48h p.i.) infected Calu-3 cells that were left untreated and/or were treated with the indicated amounts of indicated IFNs or pro-inflammatory cytokines as assessed by qPCR. Lines represent the mean of n=2±SD. **b**, Correlation between average inhibition of the indicated innate immune signaling pathway and impact on replication of SARS-CoV-2 after treatment with the respective cytokine. r, Pearson’s correlation. **c**, SARS-CoV-2 N RNA in the supernatant of SARS-CoV-2 (MOI 0.05, 48h p.i.) infected Calu-3 cells that were left untreated and/or were treated with the indicated combinations of indicated IFNs (5 U/ml) or Rapamycin (125 nM) either 24 h before the infection (Pre-treatment) or 6 h post infection (Post-treatment). Dots represent individual experiments, line the mean. Fold reduction compared to control is indicated. **d**, Immunoblot analysis of the SARS-CoV-2 infection using the WCLs of Calu-3 cells in (c). Blots were stained with anti-SARS-CoV-2 S, anti-SARS-CoV-2 N, and anti-GAPDH.

In contrast to type II and II IFN signaling, autophagic turnover was strongly repressed by at least four SARS-CoV-2 proteins (Fig. 1c and Fig. 2). Thus, based on our inhibition data (Fig. 1c) we would expect that modulation of autophagy only weakly affects SARS-CoV-2 replication. Indeed, treatment with Rapamycin, which induces autophagy, reduced viral replication to a maximum of 4-6-fold (Supplementary Fig. 5e). Bafilomycin A1, which blocks autophagy, had little to no effects (Supplementary Fig. 5e). Both drugs were used at concentrations that only marginally affected cell survival (Supplementary Fig. 5f).

Thus, our results indicate that the overall efficiency of SARS-CoV-2 proteins in counteracting specific signaling pathway is predictive for the overall antiviral potency of the pathway, as illustrated by different types of IFNs.

### Rational innate immune activation allows highly effective control of SARS-CoV-2

IFN therapy is commonly associated with significant adverse effects, due to induction of inflammation. To minimize detrimental pro-inflammatory effects of IFNs, doses required for efficient viral restriction should be reduced. Thus, we analyzed the impact of the most potent IFNs, IFN-γ and IFN-λ1 and their combination of SARS-CoV-2. To mimic prophylactic and therapeutic treatment we examined pretreatment for 24 h before infection with SARS-CoV-2 and treatment 6 h post-infection. Overall, the effects of IFN treatment were about 10-fold stronger in the prophylactic condition than in the therapeutic treatment but consistent (Fig. 5c, d). Expression analysis of SARS-CoV-2 S and N confirmed the qPCR results, and equal GAPDH levels exclude effects on viral replication by cytotoxicity (Fig. 5d). While treatment with a single dose of IFN-γ and IFN-λ1 alone reduced viral RNA production 50-100-fold, the combinatorial treatment at the same concentration potentiated the effect to about 1000-fold reduction in SARS-CoV-2 RNA (Fig. 5c).

To further decrease inflammatory side-effects by IFN treatment, anti-inflammatory pathways like autophagy could be induced^53–55^. Treatment with Rapamycin, which induces autophagy, already reduces viral replication ~ 4-6-fold at 125nM (Fig. 5c). Treatment of Rapamycin (125nM) in combination with either IFN-γ or IFN-λ1 was found to be additive (Fig. 5c, d). Triple treatment with IFN-γ, IFN-λ1 and Rapamycin showed the most potent anti-viral effect of all combinations for pre-treatment and posttreatment, reducing viral RNA in the supernatant by 2100-fold and 85-fold, respectively (Fig. 5c).

In summary, our data shows that the anti-SARS-CoV-2 effect of combinatorial treatments of IFN-γ, IFN-λ1 are synergistic. Additional activation of anti-inflammatory autophagy by Rapamycin further decreased SARS-CoV-2 replication. This suggests that concerted activation of innate immunity may be an effective anti-viral approach, exploiting vulnerabilities of SARS-CoV-2 revealed by analysis of its innate immune antagonism.

## DISCUSSION

Viruses drastically alter our innate immune defenses to establish an infection and propagate to the next host^13,14,21,27,45,56^. Our data reveal the extend of immune manipulation SARS-CoV-2 employs. We determined the major antagonists of type I IFN induction and signaling as well as pro-inflammatory NF-κB activity encoded by SARS-CoV-2 (Nsp1, Nsp5, Nsp13, Nsp14, ORF6 and ORF7b). Type II and III IFN signaling is targeted by a similar set of proteins, although much less efficient. Autophagy is majorly targeted by Nsp15, ORF3a, E, M and ORF7a. Inflammasome activity is very weakly induced by Nsp8, Nsp11 and ORF9c, but none of the SARS-CoV-2 proteins block formation of the NLR4C inflammasome. Subsequent mechanistic studies revealed that SARS-CoV-2 proteins synergistically block type-I IFN signaling and autophagy at various levels. We could reveal for the first time, that Nsp14 lowers the cellular levels of the IFN receptor, IFNAR, thus blocking activation of the crucial transcription factors STAT1 and STAT2. Both ORF3a and ORF7a cause fragmentation of the TGN via disturbing the late endosomal pathway. This is a common strategy of viruses to block autophagic turnover. Examination of the functional conservation showed that SARS-CoV-2 Nsp15 was less efficient in blocking innate immune activation, both type I IFN induction and signaling, than SARS-CoV-1 Nsp15. This may ultimately cause SARS-CoV-2 to be better controlled by the innate immune system than SARS-CoV-1, explaining higher numbers of subclinical infections and thus overall lower mortality rates of the current pandemic CoV. Overall, the combined analysis of IFN antagonism allowed us to deduce that treatment with type-I IFNs and regulation of autophagy is only weakly anti-viral. In contrast, treatment with IFN-γ and IFN-λ1 drastically reduced SARS-CoV-2 replication. Finally, combinatorial treatment of SARS-CoV-2 with these two IFNs potentiated the effects of the individual treatments. This may pave the way for future anti-viral therapies against SARS-CoV-2 based on rational innate immune activation.

Why would multiple effective proteins target the same pathway? For example, type I IFN signaling could have been shut down by Nsp1, Nsp5, Nsp13, Nsp14, ORF6 and ORF7b alone, each reducing the activation of the innate immune pathways to below 10%. However, our assays revealed (Figs. 1–3) that the targeting mechanisms are often not redundant and may act synergistically. This could allow the virus to better control the targeted pathway, thus minimizing the effect of the signaling on its replication. In addition, a viral protein majorly targeting one pathway may affect other connected immune pathways at once. For example, disturbance of the kinase TBK1 activation may affect primarily IFN induction and to a lesser extend also impact autophagy^57^. Proteome analyses revealed the late endosome/Golgi network as a target of ORF3a and ORF7a. Our data suggests, that both ORF3a and ORF7a of SARS-CoV-2 cause fragmentation of Golgi apparatus and thus blockage of autophagy. SARS-CoV-1 ORF3a was previously already implicated in Golgi fragmentation, thus our data suggests that SARS-CoV-2 ORF3a uses a similar strategy^51,58^. Notably, fragmentation of the Golgi is for example triggered by Hepatitis C virus viruses to block anti-viral autophagic turnover^50^ and thus may represent a common strategy by viruses to avoid autophagic degradation. Based on our initial proteome approaches, future studies will see more mechanistic data to explain the molecular details of the impact of SARS-CoV-2 proteins on innate immune activation. Notably, several proteins including ORF6, ORF3a, ORF7a, M and E accumulate at the Golgi network or in perinuclear spaces, alluding to the emerging role of the Golgi as a hub for immune manipulation^52,59^.

Our results demonstrate that ORF6, ORF3a, ORF7a and ORF7b are the strongest innate immune antagonists among the accessory genes of SARS-CoV-2 (Fig. 1). Besides the accessory genes, which classically encode immune antagonists, a surprising number of non-structural proteins manipulate innate immunity. Nsp1, which targets cellular translation and thus broadly inhibits any response dependent on cellular translation, which includes IFN induction and expression of ISGs and anti-viral effector proteins^30^. However, Nsp3, Nsp5, Nsp13 and to a lower extend Nsp15 also antagonized IFN induction and signaling. These non-structural proteins of CoVs have important functions in the viral life-cycle: Nsp3 as ISG/ubiquitin ligase and protease for autocatalytic processing of the ORF1a/b precursor protein^60–62^ Nsp5 as a protease mediating cleavage of the precursor polyproteins^63,64^, Nsp13 as NTPase/Helicase^65,66^ and Nsp15 as endoribonuclease^67^. It is so far not completely clear how their enzymatic functions may impact their activity against innate immunity. Except for Nsp3, as its activity as a de-ISGlase may inactivate the transcription factor IRF3 and thus reduce IFN induction^62^. According to our analysis the structural proteins E and M strongly manipulated autophagy (Fig. 1d). This suggests that the incoming virion may already block autophagic turnover to prevent their own degradation by autophagy.

However, while we may pick up most counteraction strategies, our screening approach may miss immune evasion strategies employed by SARS-CoV-2. For example, many non-structural proteins form complexes, that are not formed during single overexpression and may only be functional as a full assembly. Evasion mechanisms based on RNA structures and sequences are lost due to usage of codon-optimized expression plasmids. Finally, the virus itself may employ strategies to hide itself from recognition, thus not activating innate immune defenses in the first place. One example is the capping of its genomic and sub genomic mRNAs, which removes the free triphosphate 5’end, which otherwise would be immediately recognized by the cytoplasmic sensor RIG-I.

Our analyses further revealed that the human innate immune antagonism is largely conserved in the SARS-CoV-2 closest related bat isolate, RaTG13 (Fig. 4). This indicates that the bat virus is capable of counteracting the human immune defenses, which may have facilitated successful zoonotic transmission from bat eventually to humans. Currently, the intermediate animal host of SARS-CoV-2 is under debate^3,68–70^, however it is likely, that the virus isolated from it is even closer related to SARS-CoV-2 than RATG13. Thus, any immune evasion mechanisms conserved between SARS-CoV-2 and RATG13, is likely to be conserved in the direct progenitor virus of SARS-CoV-2. The previous epidemic and related human SARS-CoV-1 and the current pandemic SARS-CoV-2 differ in susceptibility towards IFN s with SARS-CoV-1 being more resistant^26^. Furthermore, infection with SARS-CoV-2 is often asymptomatic and likely controlled by the host^26^ as lower mortality rates and higher subclinical infections suggest^4^. Paradoxically, this may support the fast spread of the virus. Thus, SARS-CoV-2 may have found the ‘perfect’ balance. Intermediate immune evasion and thus intermediate pathogenicity to support spread, but not kill the host. Our data shows that SARS-CoV-2 Nsp15 is strikingly less in efficient in IFN evasion than Nsp15 of SARS-CoV. These data are the first mechanistic evidence why SARS-CoV-1 is less susceptible towards IFN treatment than SARS-CoV-2. It may be tempting to speculate that common cold CoVs counteract the innate immune system less efficiently than SARS-CoV-2.

Our analysis indicates that during a SARS-CoV-2 infection less cytokines than expected are released, autophagic turnover is blocked and general immune activation is perturbed. This is supported by a large amount of data from COVID19 patients^24–28,45,62,71–73^. However, an important question remains: Why are some innate immune pathways, such as IFN-γ signaling less antagonized (Fig. 1)? Are the viral immune manipulation strategies ineffective? Indeed, IFN-γ is most active against SARS-CoV-2 among the IFNs^72^ (Fig. 5). One possible explanation would be that there was no need for the virus to antagonize them. Indeed, in COVID19 patients and in vitro infections with SARS-CoV-2, IFN-γ levels are surprisingly low^28,73^. Furthermore, despite high IFN-γ levels being a hallmark of cytokine storms induced by influenza viruses, the SARS-CoV-2 cytokine storm only has low IFN-γ levels and decreased IFN-γ expression in CD4+ T cells is associated with severe COVID19^4,74,75^. It is tempting to speculate that T-cells which confer pre-existing immunity against SARS-CoV-2^76,77^ could, upon activation, release IFN-γ, whose innate immune signaling may also contribute to increased clearance of the infection. Strikingly, our work thus shows that analysis of the innate antagonism may be predictive for therapeutic opportunities.

Severe side effects are prevalent for treatments with IFNs^35–37^. However, the side-effects are dose-dependent^78^. Thus, minimizing the dose required for treatment is paramount. Our data indicates that effects of treatment with multiple IFNs is additive but synergistic and potentiates each other (Fig. 5). Thus, a promising anti-viral approach may be a combinatorial treatment of different cytokines, effectively also reducing the burden of side-effects. The side effects of IFN therapy are mainly caused by inflammation. Combined with anti-inflammatory approaches such as autophagy activation by Rapamycin^54,55^, this approach may even be more successful, as our *in vitro* data suggests. Future studies are highly warranted to study rational, concerted innate immune activation against SARS-CoV-2 *in vivo.* These studies may eventually pave the way for novel therapies, which may not only work against SARS-CoV-2, but also against other pathogenic viruses, including potentially future CoVs.

In summary, our results reveal the extend of innate immune manipulation of SARS-CoV-2. Comparison to SARS-CoV-1 revealed that mutations in Nsp15 may be responsible for the higher susceptibility of SARS-CoV-2 against IFNs. Finally, our data allowed us to deduce a potent immune activation strategy against SARS-CoV-2: combinatorial application of IFN-γ and IFN-λ.

## AUTHOR CONTRIBUTIONS

L.K., M.Hi., M.Ha. performed the majority of the experimental work with help from J.H.S.. R.N. performed experiments with infectious SARS-CoV-2 assisted by F.Z.. C.B.P. performed additional experimental work. C.M.S. and S.S. generated expression constructs. J.A.M. and C.C. performed the SARS-CoV-2 infection for the proteome analysis. F.I.S. and M.C. provided the inflammasome analysis. A.I., I.F. and W.A performed the proteome analyses and the bioinformatic interrogation of the data. J. M., D.S., A.I., S.S. and K-K.C. provided resources and comments for the manuscript. K.M.J.S and F.K. conceived the study, planned experiments and wrote the manuscript. All authors reviewed and approved the manuscript.

## ACKNOWLEDGEMENTS

We thank Regina Burger, Susanne Engelhart, Daniela Krnavek, Kerstin Regensburger, Martha Meyer, Birgit Ott and Nicola Schrott for excellent technical assistance. We would like to especially acknowledge the library of SARS-CoV-2 expression plasmids which were generously given to us by Nevan Krogan (University of California, San Francisco). Additionally, we thank Dorota Kmiec (Kings College, London) for critical comments and input. This study was supported by DFG grants to F.K., J.M., K.M.J.S., D.Sa., F.I.S., A.I. and KKC (CRC1279, SPP1923, SP1600/4-1, CRC1309, Project-ID 369799452 – TRR237), EU’s Horizon 2020 research and innovation program to J.M. (Fight-nCoV, 101003555), a COVID-19 research grant of the Federal Ministry of Education and Research (MWK) Baden-Württemberg (to D.S. and F.K.) as well as the BMBF to F.K., D.Sa. and K.M.J.S. (Restrict SARS-CoV-2, protACT and IMMUNOMOD).

## DATA AVAILABILITY

Mass spectrometry datasets generated during and/or analyzed during the current study are available in the PRIDE partner repository with the dataset identifier PXD021899. The datasets generated during and/or analyzed during the current study are either included in the study and/or available from the corresponding author on reasonable request.

## MATERIAL AND METHODS

### Cell lines and cell culture and viruses

HEK293T cells were purchased from American type culture collection (ATCC: #CRL3216). The construction of HEK293T GL cells and HeLa GL cells was reported previously^42^. These cell lines were cultivated in Dulbecco’s Modified Eagle Medium (DMEM, Gibco) supplemented with 10% (v/v) fetal bovine serum (FBS, Gibco), 100 U/ml penicillin (PAN-Biotech), 100 μg/ml streptomycin (PAN-Biotech), and 2 mM L-glutamine (PANBiotech). Calu-3 (human epithelial lung adenocarcinoma, kindly provided and verified by Prof. Frick, Ulm University) cells were cultured in Minimum Essential Medium Eagle (MEM, Sigma) supplemented with 10% (v/v) FBS (Gibco) (during viral infection) or 20% (v/v) FBS (Gibco) (during all other times), 100 U/ml penicillin (PAN-Biotech), 100 μg/ml streptomycin (PAN-Biotech), 1 mM sodium pyruvate (Gibco), and 1x non-essential amino acids (Gibco). Vero E6 (Cercopithecus aethiops derived epithelial kidney cells, ATCC) cells were grown in Dulbecco’s modified Eagle’s medium (DMEM, Gibco) which was supplemented with 2.5% (v/v) fetal bovine serum (FBS, Gibco), 100 U/ml penicillin (PANBiotech), 100 μg/ml streptomycin (PAN-Biotech), 2 mM L-glutamine (PANBiotech), 1 mM sodium pyruvate (Gibco), and 1x non-essential amino acids (Gibco). All cells were cultured at 37°C in a 5% CO2, 90% humidity atmosphere. Sendai Virus was a kind gift from Prof. Hans-Georg Koch, Institute for Biochemistry and Molecular Biology, University of Freiburg. Viral isolates BetaCoV/France/IDF0372/2020 (#014V-03890) and BetaCoV/Netherlands/01/NL/2020 (#010V-03903) were obtained through the European Virus Archive global.

### Expression constructs and plasmids

pLVX-EF1alpha constructs containing all strep II tagged, codon optimized open reading frames (Orfs) of SARS-CoV-2 (control, Nsp1, Nsp2, Nsp3, Nsp4, Nsp5, Nsp6, Nsp7, Nsp8, Nsp9, Nsp10, Nsp11, Nsp12, Nsp13, Nsp14, Nsp15, Nsp16, S, ORF3a, ORF3c, E, M, ORF6, ORF7a, ORF7b, ORF8, N, ORF9b, ORF9c, and ORF10) were a kind gift by David Gordon and Nevan Krogan. V5 tagged, codon optimized Orfs coding for Nsp1, Nsp3, Nsp7, Nsp15, M, N, Orf3a, Orf6, and Orf7a from SARS-CoV-2, RaTG13, and SARS-CoV-1 were synthesized by Twist Bioscience and subcloned into the pCG vector using restriction cloning using the restriction enzymes XbaI and MluI (New England Biolabs). Firefly luciferase reporter constructs, harboring binding sites for NF-κB or IRF3, ISRE or GAS sites, or the genomic promoter of IFNA4 or IFNB1 in front of the reporter were previously described^79,80^. The GAPDH_PROM_01 Renilla SP Luciferase construct was purchased from switchgear genomics. pCR3 constructs coding for FLAG-tagged Measles morbillivirus V (MeV V) protein or Rabies virus P (RABV P) protein were described previously^79,81^. pEGFP-N1_hTRIM3 2^82^ was a gift from Martin Dorf (Addgene, #69541), the Orf of TRIM32 was subcloned into the pIRES_FLAG vector using Gibson assembly as previously described^42^. The pCMV6 construct coding for myc-FLAG tagged GNG5 was purchased from OriGene.

### Transfections

Plasmid DNA was transfected using either the TransIT-LT1 Transfection Reagent (Mirus) or Polyethylenimine (PEI, 1 mg/ml in H2O, Sigma-Aldrich) according to the manufacturers recommendations or as described previously^42,83^.

### Luciferase assays

HEK293T cells were transiently transfected with firefly luciferase reporter constructs, renilla luciferase control constructs, and constructs expressing SARS-CoV-2 Orfs in 48-well plates using TransIT-LT1. One day post-transfection, the cells were stimulated with IFN-β (1,000 U/ml, 8 h, Merck), IFN-α2 (500 U/ml, 24 h, Sigma-Aldrich), IFN-γ (400 U/ml, 24 h, Sigma-Aldrich), IFN-λ1 (100 ng/ml, 8 h, R&D Systems), IL-1α (10 ng/ml, 24 h, R&D Systems), TNFα (25 ng/ml, 24 h, Sigma-Aldrich), or SeV (1:500, 24 h, kindly provided by Hans-Georg Koch, Freiburg). 8-24 h post-stimulation, the cells were lysed in passive lysis buffer and luciferase activities of the firefly luciferase and renilla luciferase were determined using the dual-glo luciferase assay system (Promega) and an Orion II microplate Luminometer (Berthold). Cell viability of the transfected cells was measured using the CellTiter-Glo Luminescent Cell Viability Assay (Promega).

### Cell viability assay

Calu-3 or HEK293T cells were treated with cytokines or transiently transfected using TransIT-LT1. To measure metabolic activity, cells were lysed in passive lysis buffer and analyzed using the CellTiter-Glo Luminescent Cell Viability Assay (Promega) according to manufacturer’s instructions and an Orion II microplate Luminometer (Berthold).

### Autophagy quantification by flow cytometry

The number of Autophagosomes was quantified as previously described^42^, either in a basal state, or stimulated with rapamycin (1μM, Sigma) or Bafilomycin A1 (0.1 μM, Santa Cruz Biotechnology). In brief, HEK293T cells stably expressing GFP-LC3B (HEK293T GL) were transiently transfected using PEI. 48 h post-transfection, cells were harvested in PBS and treated for 20 min at 4 °C with PBS containing 0.05% Saponin. Non-membrane bound GFP-LC3B was washed out of the permeabilized cells using PBS twice, followed by fixation in 4% Paraformaldehyde (Santa Cruz Biotechnology). The fluorescence intensity of membrane associated GFP-LC3B was then quantified via flow cytometry (FACSCanto II, BD Biosciences). The GFP-LC3B mean fluorescence intensity of the control (baseline) was subtracted.

### Generation of stable THP-1 cells

THP-1 cell liens were generated by transduction with lentivirus generated with the indicated pLVX-EF1alpha vectors (kind gift from Nevan Krogan), as well as packaging vectors psPax2 and pMD2.G (kind gifts from from Didier Trono (Addgene plasmid #12259 and 12260).

### Inflammasome activity quantification

To quantify NLRC4 inflammasomes, THP-1 EGFP inflammasome reporter cells (to be described elsewhere) were differentiated in 50 ng/mL PMA for 16 h, followed by induction of gene expression with 1 ug/mL doxycycline for 24 h. Cells were subsequently treated with 1 ug/mL Bacillus anthracis PA and 0.1 ug/mL LFn-MxiH for 1h in the presence of 40 uM Vx-765. Cells were trypsinized, fixed in formaldehyde, and analyzed by flow cytometry. Cells with ASC specks were defined exploiting the characteristic distribution of EGFP in height versus width plots as described previously^84^.

### Whole-cell lysates

Whole-cell lysates were prepared by collecting cells in Phosphate-Buffered Saline (PBS, Gibco). The cell pellet (500 g, 4 °C, 5 min) was lysed in transmembrane lysis buffer [50 mM HEPES pH 7.4, 150 mM NaCl, 1% Triton X-100, 5 mM ethylenediaminetetraacetic acid (EDTA)] by vortexing at maximum speed for 30 s. Cell debris were pelleted by centrifugation (20,000 g, 4°C, 20 min) and the total protein concentration of the cleared lysates was measured using the Pierce BCA Protein Assay Kit (Thermo Scientific) according to manufacturer’s instructions. The lysates were adjusted to the same protein concentration and stored at −20°C.

### SDS-PAGE and immunoblotting

SDS-PAGE and immunoblotting was performed using standard techniques as previously described^42^. In brief, whole cell lysates were mixed with 6x Protein Sample Loading Buffer (LI-COR, at a final dilution of 1x) supplemented with 15% β-mercaptoethanol (Sigma Aldrich), heated to 95°C for 5 min, separated on NuPAGE 4-12% Bis-Tris Gels (Invitrogen) for 90 minutes at 100 V and blotted onto Immobilon-FL PVDF membranes (Merck Millipore). The transfer was performed at a constant voltage of 30 V for 30 min. After the transfer, the membrane was blocked in 1% Casein in PBS (Thermo Scientific). Proteins were stained using primary antibodies against β-actin (1:10,000, AC-15, Sigma), strep II-tag (1:1,000, NBP2-43735, Novus), strep II-tag (1:2,000, ab76949, abcam), GAPDH (1:1,000, 607902, Biolegend), pSTAT1 (1:1,000, Y701, Cell Signaling Technology), STAT1 (1:1,000, 9172S, Cell Signaling Technology), pSTAT2 (1:1,000, Y690, Cell Signaling Technology), STAT2 (1:1,000, 4594S, Cell Signaling Technology), IFNAR1 (1:1,000, ab45172, abcam), p62 (1:1,000, GP62-N, ProGen), LC3α/β (1:200, G-4, Santa Cruz Biotechnology), Beclin-1 (1:1,000, 3738S, Cell Signaling Technology), ULK1 (1:1,000, D8H5, Cell Signaling Technology), Rab5 (1:1,000, C8B1, Cell Signaling Technology), SARS-CoV-2 Nsp3 (1:1,000, GTX135614, GeneTex), Flag-tag (1:5,000, M2, Sigma), V5-tag (1:1,000, D3H8Q, Cell Signaling Technology), SARS-CoV-2 (COVID-19) spike antibody (1:1000, 1A9, Biozol), SARS-CoV/SARS-CoV-2 Nucleocapsid Antibody (1:1000, MM05, SinoBiological), and Infrared Dye labelled secondary antibodies (1:20,000, LI-COR IRDye), diluted in 0.05% Casein in PBS. Band intensities were quantified using Image Studio (LI-COR) and protein levels were normalized on β-actin or GAPDH levels.

### Immunofluorescence

HeLa GL cells were transfected using TransIT-LT1 and grown on coverslips in 24-well plates. The cells were fixed using 4% PFA (Santa Cruz Biotechnology), and permeabilized and blocked with PBS containing 0.5% Triton X-100 (Sigma) and 5% FBS (Gibco). The cells were stained using primary antibodies against strep II-tag (1:200, NBP2-43735, Novus) and TGN46 (1:400, AHP500GT, Bio Rad), secondary antibodies fluorescently labelled with AlexaFluor568 targeting rabbit-IgGs (1:400, A10042, Invitrogen) and AlexaFluor647 targeting sheep-IgG (1:400, A21448, Invitrogen), and DAPI (1:1,000) to stain nuclei. The coverslips were mounted on microscope slides using mowiol mounting medium (10% (w/v) Mowiol 4-88, 25% (w/v) Glycerol, 25% (v/v) water, 50% (v/v) Tris-Cl 0.2M pH 8.5, 2.5% (w/v) DABCO). Images were acquired using a Zeiss LSM710 and analyzed with Fiji ImageJ.

### Autophagy quantification by counting

HeLa GL cells were transfected using TransIT-LT1 and grown on coverslips in 24-well plates. The cells were treated and stained for the transfected proteins as described in the Immunofluorescence method-paragraph. After acquiring images of 30+ transfected cells, the total pixel area of GFP-LC3B puncta per cell was quantified using Fiji ImageJ as previously described^42^. In brief, the channels were separated to work with the GFP-channel, the background removed and smoothed, and a threshold was applied to isolate the GFP-LC3B puncta. By analysing the particles, the total area was determined. Cells treated with 1 μM chloroquine overnight were used as positive control.

### RT-qPCR

SARS-CoV-2 N (nucleoprotein) transcript levels were determined as previously described^72,83^. In brief, supernatants were collected from SARS-CoV-2 infected Calu-3 cells 48 h post-infection. Total RNA was isolated using the Viral RNA Mini Kit (Qiagen, Cat# 52906) according to the manufacturer’s instructions. RT-qPCR was performed as previously described using TaqMan Fast Virus 1-Step Master Mix (Thermo Fisher, Cat#4444436) and an OneStepPlus Real-Time PCR System (96-well format, fast mode). Primers were purchased from Biomers (Ulm, Germany) and dissolved in RNase free water. Synthetic SARS-CoV-2-RNA (Twist Bioscience) or RNA isolated from BetaCoV/France/IDF0372/2020 viral stocks quantified via this synthetic RNA (for low Ct samples) were used as a quantitative standard to obtain viral copy numbers. (Forward primer (HKU-NF): 5’-TAA TCA GAC AAG GAA CTG ATT A-3’; Reverse primer (HKU-NR): 5’-CGA AGG TGT GAC TTC CAT G-3’; Probe (HKU-NP): 5’-FAM-GCA AAT TGT GCA ATT TGC GG-TAMRA-3’.

### Inhibition of SARS-CoV-2 by immune modulation

300,000 Calu-3 cells were seeded in 12-well plates. The cells were stimulated with increasing amounts of IFNs (α2, β and γ, 0.8, 4, 20, 100 and 500 U/ml or λ1, 0.16, 0.8, 4, 20 and 100 ng/ml) at 24 h and72 h post-seeding, with an intermediate medium change at 48 h post-seeding. 2 h after the second stimulation, the cells were infected with SARS-CoV-2 (MOI 0.05) and 6 h later the medium was changed. 48 h postinfection, the cells were harvested for further analysis.

### Propagation of SARS-CoV-2

BetaCoV/Netherlands/01/NL/2020 and BetaCoV/ France/IDF0372/2020 were obtained from the European Virus Archive. The viruses were propagated by infecting 70% confluent Vero E6 in 75 cm^2^ cell culture flasks at a MOI of 0.003 in 3.5 ml serum-free medium containing 1 μg/ml trypsin. The cells were then incubated for 2 h at 37°C, before adding 20 ml medium containing 15 mM HEPES. Three days post-infection, the medium was exchanged and the supernatants were harvested 5 days post-infection upon visible cytopathic effect. The supernatants were cleared by centrifugation, aliquoted and stored at −80°C. The infectious virus titre was determined as plaque forming units (PFU).

### Proteome analysis

For the proteome analysis of infected cells, 0.6×106 Caco-2 cells were infected with SARS-CoV-2 BetaCoV/Netherlands/01/NL/2020 at an MOI of 0.5 and harvested 24 h and 48 h post infection with TM lysis buffer supplemented with 1:500 protease inhibitor. After centrifugation for 10 minutes with 14,000 rpm at 4°C, supernatant was mixed with 6xLaemmli buffer and stored at −20°C until further analysis. Then, the samples were boiled at 95°C for 10 minutes to ensure denaturation. For the proteome analysis of single overexpressed SARS-CoV-2 proteins, 1×107 HEK293T cells were transfected with the respective constructs (pCG vectors containing V5 tagged, codon optimized open reading frames (Orfs) of SARS-CoV-2 (Nsp1, Nsp7, Nsp15, Nsp16, S, E, M, N, ORF3a, ORF6, ORF7a)). The cells were harvested in PBS and processed for LC-MS using the iST-kit (Preomics) as recommended by the manufacturer. For LC-MS purposes, desalted peptides were injected in a nanoElute system (Bruker) and separated in a 25-cm analytical column (75μm ID, 1.6μm C18, IonOpticks) with a 100-min gradient from 2 to 37% acetonitrile in 0.1% formic acid. The effluent from the HPLC was directly electrosprayed into a hybrid trapped ion mobility-quadrupole time-of-flight mass spectrometer (timsTOF Pro, Bruker Daltonics, Bremen, Germany) using the nano-electrospray ion source at 1.4kV (Captive Spray, Bruker Daltonics). The timsTOF was operated at 100% duty cycle in data dependent mode to automatically switch between one full TIMS-MS scan and ten PASEF MS/MS scans in the range from 100–1700 m/z in positive electrospray mode with an overall acquisition cycle of 1.23 s. The ion mobility was scanned from 0.6 to 1.60Vs/cm2 with TIMS ion charge control set to 5e4, RF potential of 300 Vpp. The TIMS dimension was calibrated linearly using four selected ions from the Agilent ESI LC/MS tuning mix [m/z, 1/K0: (322.0481, 0.7318 Vscm-2), (622.0289, 0.9848 Vs/cm2), (922.0097, 1.1895 Vs/cm2), (1221.9906, 1.3820 Vs/cm2)]. The mass spectrometry proteomics data have been deposited to the ProteomeXchange Consortium via the PRIDE partner repository with the dataset identifier PXD021899. MaxQuant 1.6.15.0 was used to identify proteins and quantify by LFQ with the following parameters: Database, Uniprot_AUP000005640_Hsapiens_20200120.fasta supplemented with the sequences of NSP1_V5, NSP7_V5, NSP15_V5, NSP16_V5, E_V5, M_V5, N_V5, S_V5, ORF3_V5, ORF6_V5, ORF7_V5 and Spike protein from SARSCoV2^39^; MS tol, 10ppm; MS/MS tol, 20ppm Da; Peptide FDR, 0.1; Protein FDR, 0.01 Min. peptide Length, 7; Variable modifications, Oxidation (M); Fixed modifications, Carbamidomethyl (C); Peptides for protein quantitation, razor and unique; Min. peptides, 1; Min. ratio count, 2. Identified proteins were considered as differential if their MaxQuant LFQ values. Raw data was analyzed using R. Outliers (below 0.05 and above 0.95) appearing in more than 2 cases were removed as artefacts of the overexpression. Heatmaps were generated using R, using the inbuilt hierarchical clustering of heatmap.2 and displayed in Corel Draw.

### GO Analysis

From the proteome of the respective samples, proteins regulated more than 4-fold compared to the vector control were extracted and submitted to PANTHER (cellular component analysis).

### Half-life analysis

We focused the half-life comparisons to proteins for which we identified peptides that resided within the first 50 N-terminal amino acids. To do this we extracted peptides for both NSP1+ (NSP over expression) and WT (wild type) samples that fall within the first 50 AA window starting at the N-terminus from the result file (peptide.txt, Maxquant 1.6.15.0). These peptides were then mapped to the corresponding protein intensities and the relative changes of log2 transformed iBAQ values calculated and grouped into three groups: I. enriched in NSP1+: log2(fc) > 2, II. enriched in WT: log_2_(fc) < −2, III. Not enriched: −2<= log_2_(fc) <= 2. The proteins for which data on the the half lives in hepatocytes^47^ were extracted and plotted by scaling their mean half-lives corresponding to the proteins in each group to the interval [0-1] using min-max normalization and generated boxplots for each of them, which is depicted in supplementary figure 4J. We used MATLAB 2019b for the half-life analysis.

### Statistical analysis

Statistical analyses were performed using GraphPad PRISM 8 (GraphPad Software). P-values were determined using a two-tailed Student’s t test with Welch’s correction. Unless otherwise stated, data are shown as the mean of at least three biological replicates ± SEM. Significant differences are indicated as: *, p < 0.05; **, p < 0.01; ***, p < 0. 001. Statistical parameters are specified in the figure legends.

