## Supplementary Figures 1-5 for "Imperfect innate immune antagonism renders SARS-CoV-2 vulnerable towards IFN-γ and -λ"

Affiliations:

1 Institute of Molecular Virology
Ulm University Medical Center
89081 Ulm, Germany

2 Gene Center & Max von Pettenkofer-Institute of Virology
Ludwig-Maximilians-Universität München
81377 Munich, Germany

3 Biomedical Center, Zentrallabor für Proteinanalytik (Protein Analysis Unit)
Department of Molecular Biology,
Ludwig-Maximilians-Universität München
82152 Planegg-Martinsried, Germany

4 Graduate School for Quantitative Biosciences (QBM)
Ludwig-Maximilians-University of Munich,
81377 Munich, Germany

5 Institute for Medical Microbiology and Hygiene
Ulm University Medical Center
89081 Ulm, Germany

6 Institute of Innate Immunity
University of Bonn
53127 Bonn, Germany

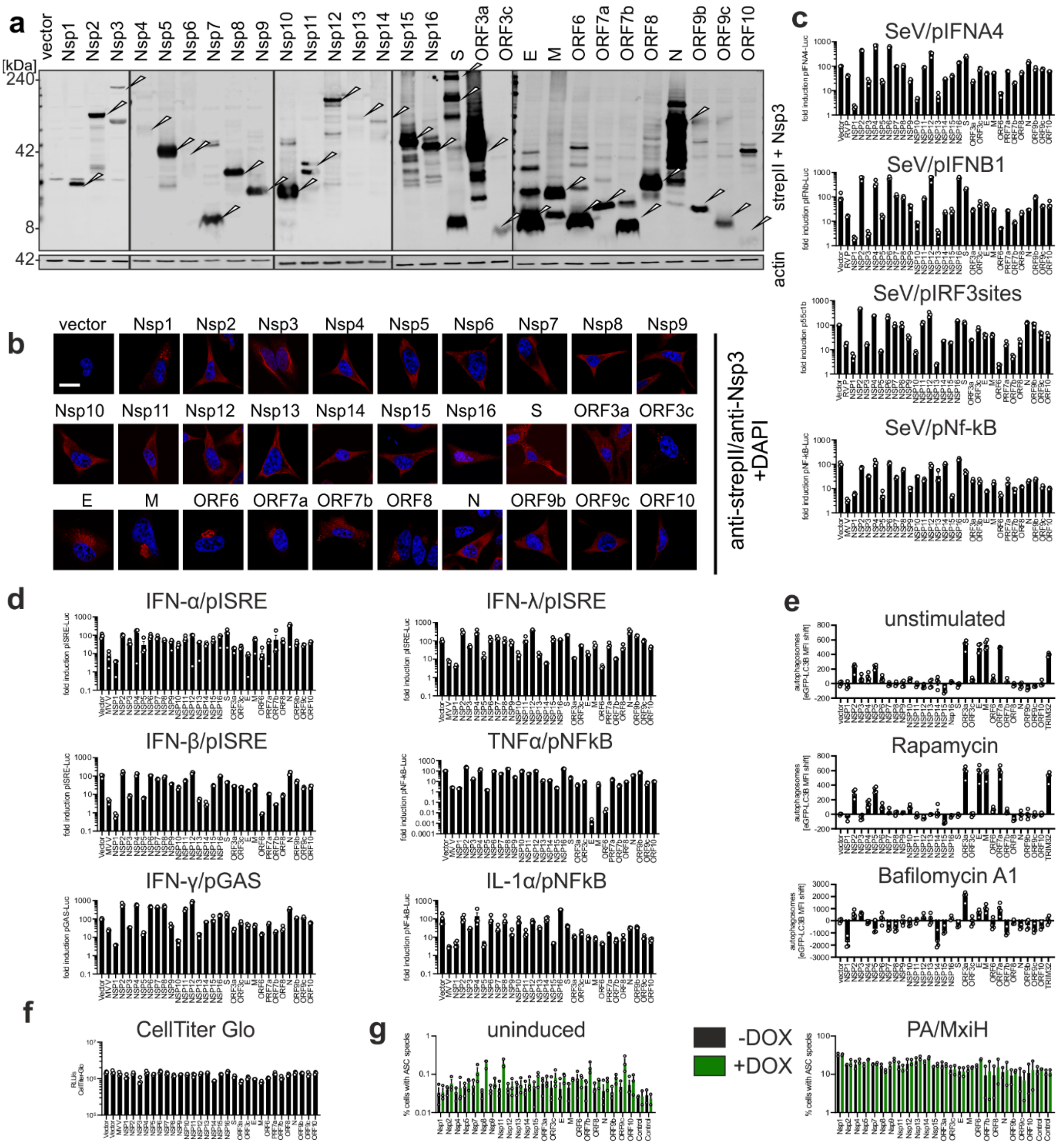


**Supplementary Figure 1: Expression and impact of 29 SARS-CoV-2 proteins on innate immune activation.** **a,** Immunoblots of whole cell lysates (WCLs) of HEK293T cells transiently transfected with the indicated SARS-CoV-2 expression plasmids stained with anti-strep II and anti-actin antibodies. **b**, Intracellular localisation of indicated SARS-CoV-2 proteins. Exemplary confocal microscopy images of Hela cells expressing the indicated 2xstrep II-tagged SARS-CoV-2 proteins (red). Nuclei, DAPI (blue). Scale bar, 10 µm. **c,** Effect of SARS-CoV-2 proteins on RLR signalling. Transiently transfected HEK293T cells expressing the indicated SARS-CoV-2 proteins as well as different Firefly luciferase promotor constructs were infected with Sendai virus or left uninfected. 24 h p.i., Firefly luciferase activities were quantified as relative light units (RLU/s). Shown are mean values of n=3±SEM, stimulated vector control set to 100%. **d**, Effect of SARS-CoV-2 proteins on interferon induction. Luciferase reporter gene assay in HEK293T cells transiently transfected with the indicates SARS-CoV-2 expression plasmids, a GAPDH promoter-Renilla luciferase construct and the indicated Firefly luciferase promotor plasmids. 24 h post transfection cells were stimulated with various interferons and cytokines. Firefly luciferase activities were quantified as relative light units (RLU/s) 8-24 h post stimulation and normalized to cell metabolic activity (CellTiter Glo). Show are mean values of n=3±SEM, stimulated vector control set to 100%. **e**, Autophagy levels in presence of SARS-CoV-2 proteins. HEK293T cells stably expressing GFP-LC3B were transiently transfected with indicated SARS-CoV-2 plasmids. Cells were stimulated with the indicated compounds or left untreated. Autophagosomes were quantified by flow cytometry as mean fluorescence intensity of GFP-LC3B-positive vesicles in saponin-permeabilised cells. Bars represent the mean of n= 4±SEM. **f,** Cell metabolic activity of HEK293T cells transfected and stimulated as described in (f) measured using the CellTiter Glo Assay Kit (Promega) according to the manufacturer's recommendations. Bars represent mean values of n=3±SEM. **g**, Percentage of ASC-GFP speckles in doxycycline (DOX) inducible THP-1 cells, expressing indicated proteins upon stimulation. Left panel, unstimulated, right panel, stimulated with PA and MxiH. Bars represent mean values of n =2-3±SEM.

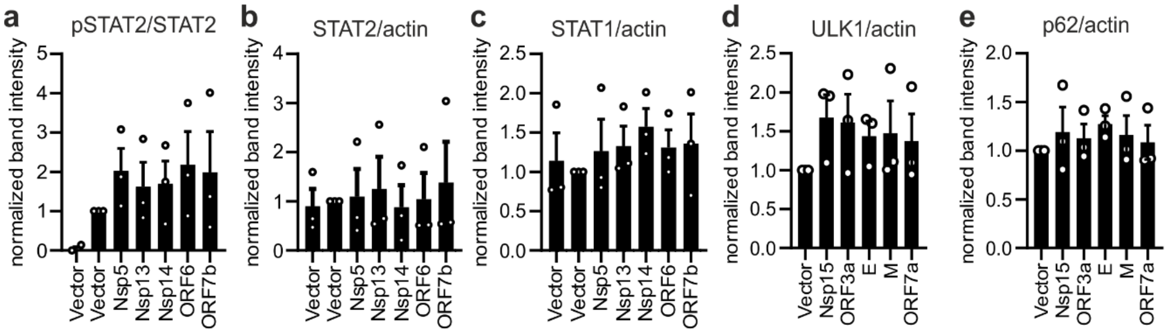


**Supplementary Figure 2: Quantification of the IFN and autophagy response in Figure 2.** Quantification of the band intensities, normalised as indicated. **a-c**, Whole cell lysates (WCLs) of HEK293T cells expressing the indicated SARS-CoV-2 proteins were stimulated with IFN-β (45 min). **d-e**, Whole cell lysates (WCLs) of HEK293T cells expressing the indicated SARS-CoV-2 proteins. Blots were stained with (a) anti-pSTAT2, anti-STAT2 (b) anti-STAT2 and anti-actin (c) anti-STAT1 and anti-actin (d) anti-ULK1 and anti-actin and (e) anti-Beclin-1 and anti-actin. Ratios were calculated as indicated on the y-axis and normalized to the stimulated vector control. Shown are mean values of n=3± SEM.

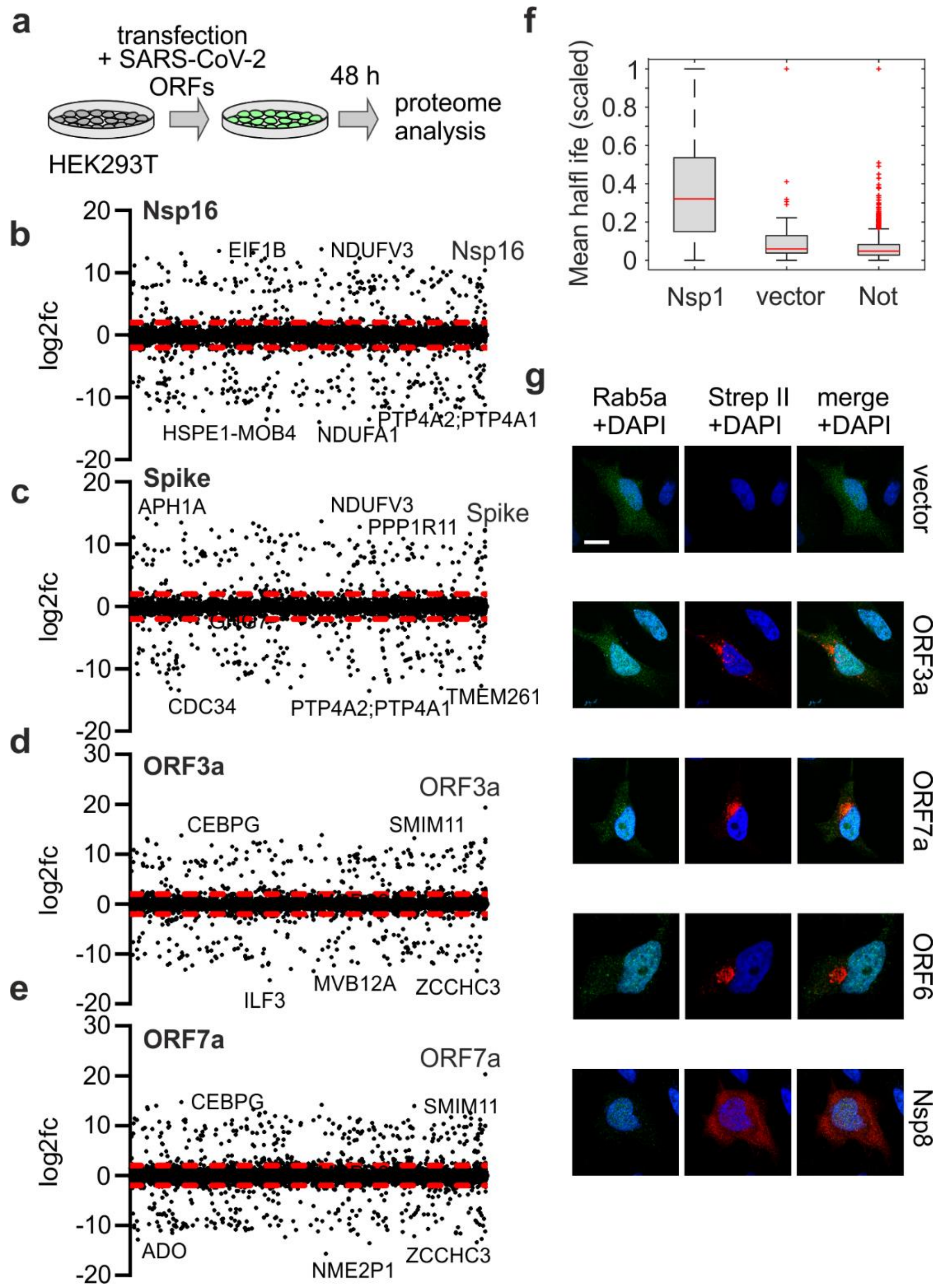


**Supplementary Figure 3: Proteome analysis and localisation of ORF3a and ORF7a. a,** Schematic depiction illustrating transient transfection of HEK293T cells with SARS-CoV-2 expression constructs for proteome analysis. **b – e**, Scatter plots of log2 fold changes in expression of proteins in HEK293T cells expressing the indicated SARS-CoV-2 ORFs. Red lines, 4-fold changes. **f**, Box and whisker plot showing the half-life of the proteins regulated by overexpression of Nsp1 or the respective controls. **g**, Exemplary confocal laser scanning microscopy images of HeLa cells transiently transfected with FLAG-Rab5α and the indicated strep II-tagged viral proteins. Cells were stained with anti-FLAG(green), anti-strep II(red). DAPI, nuclei(blue). Scale bar, 10 µM.

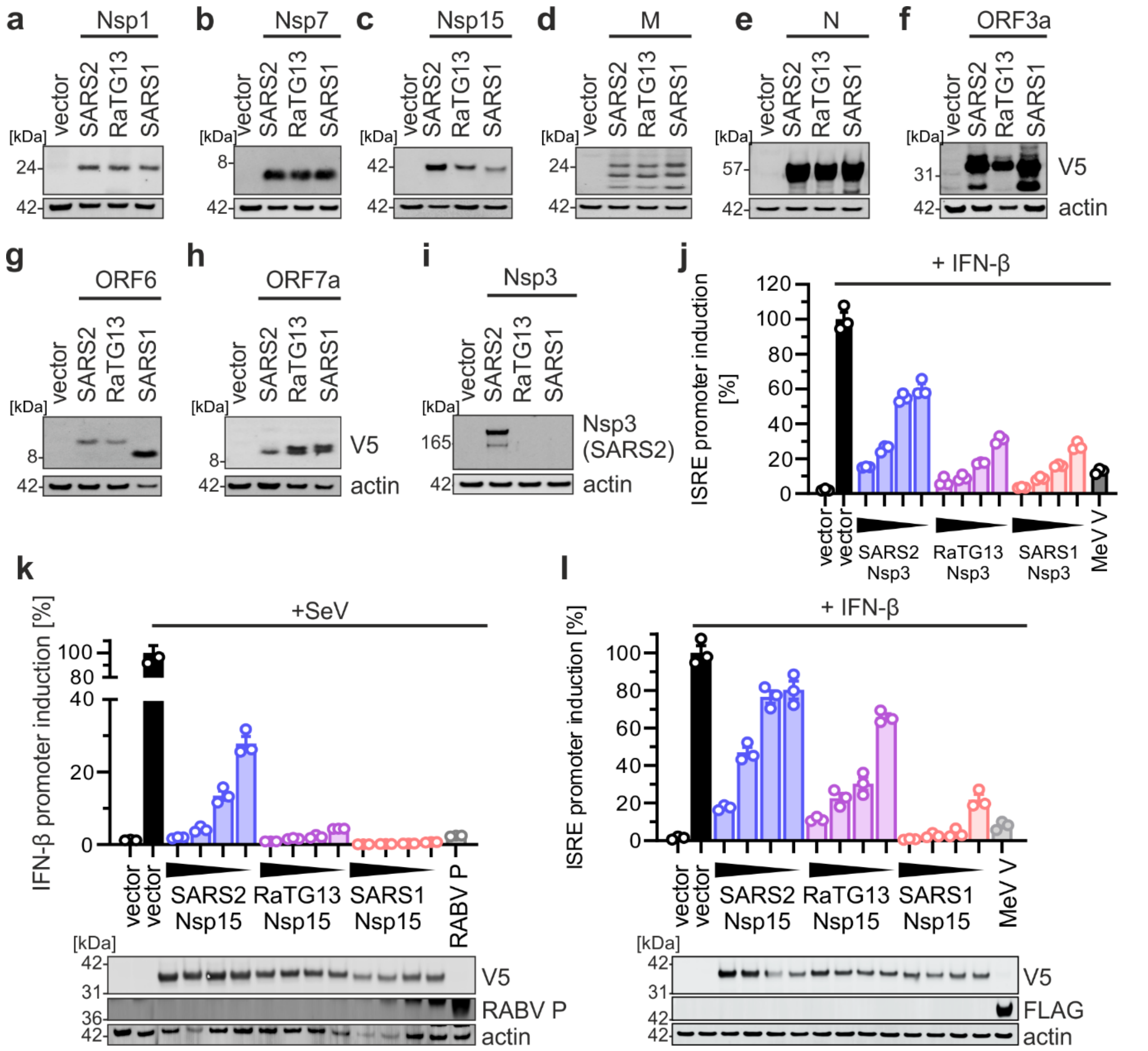


**Supplementary Figure 4: Expression of SARS-CoV-2, RaTG-13-CoV and SARS-CoV constructs and functional comparison. a – i,** Immunoblots of whole cell lysates (WCLs) of HEK293T cells transiently transfected with the indicated SARS-CoV-2, RaTG-13-CoV or SARS-CoV expression plasmids and stained with anti-V5 (a-h) and anti-GAPDH antibodies or anti-SARS-CoV-2 Nsp3 (i). **j,** pISRE Firefly luciferase promotor induction in IFN-β stimulated HEK293T cells previously transfected with the indicated amounts of V5-tagged NSP3 of SARS-CoV-2, RaTG-13-CoV or SARS-CoV. Firefly luciferase activities were quantified as RLU/s and normalized to cell metabolic activity (CellTiter Glo); stimulated vector control was set to 100%. FLAG-tagged MeV V was used as a control. Bars represent the mean values of n=3±SEM. **k**, Comparison of IFN-β antagonism of the V5-tagged NSP15 protein of SARS-CoV-2, RaTG-13-CoV and SARS-CoV as described in (j). RABV P was used as a control. Immunoblots of WCLs stained with anti-V5, anti-RABV P, anti-actin. **l**, Comparison of IFN-β antagonism of the V5-tagged NSP15 protein of SARS-CoV-2, RaTG-13-CoV and SARS-CoV as described in (j). FLAG-tagged MeV V was used as a control. Immunoblots of WCLs stained with anti-V5, anti-FLAG, anti-actin.

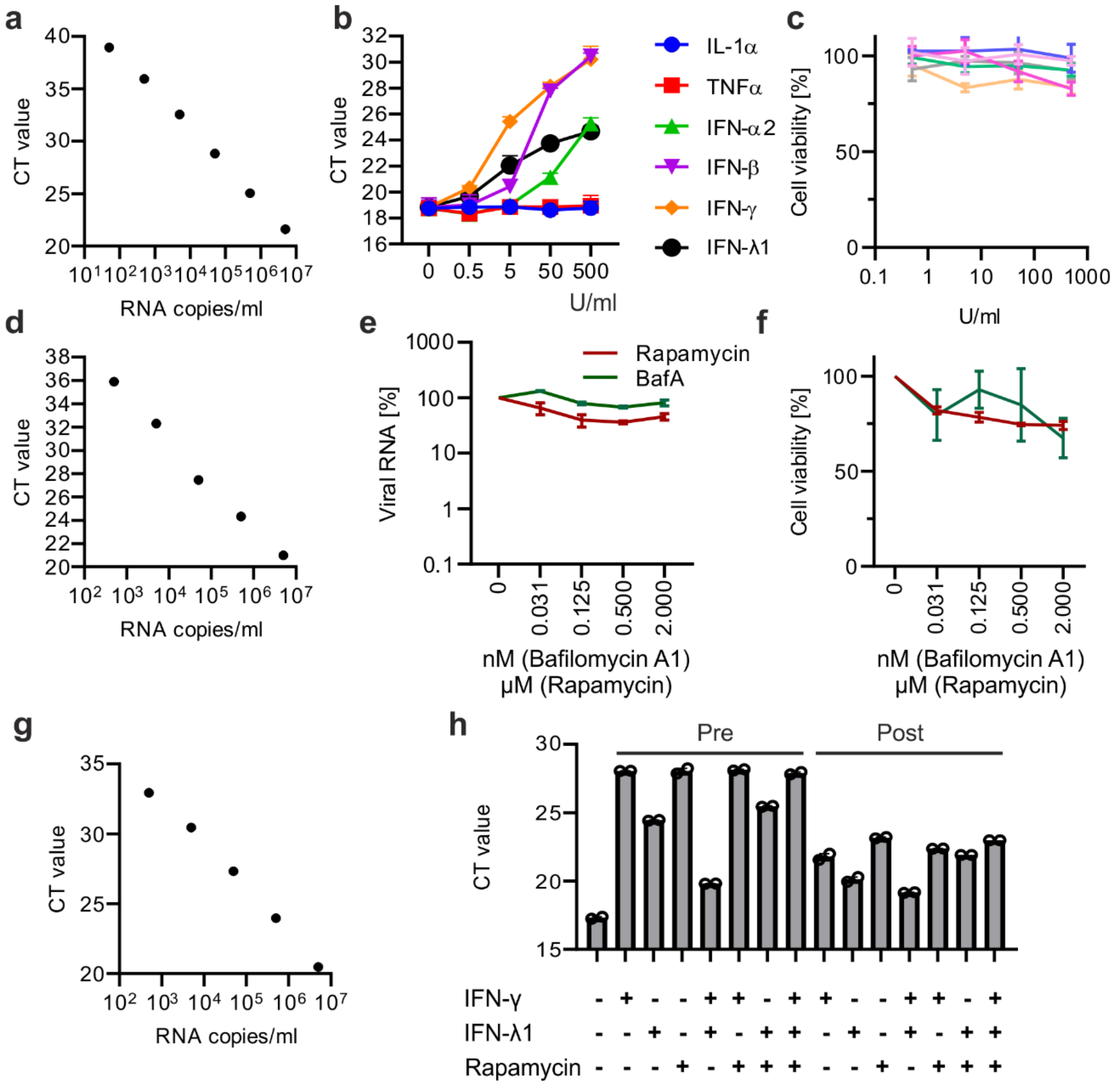


**Supplementary Figure 5: Impact of IFN and autophagy activation on SARS-CoV-2 replication. a,** Exemplary standard curve of raw qRT-PCR CT values corresponding to the SARS-CoV-2 N RNA copy numbers per ml supernatant for one replicate. **b,** Effect of indicated IFNs or pro-inflammatory cytokines in different concentrations on SARS-CoV-2 replication in Calu-3 cells. Shown are raw CT values of one replicate, n=3(technical replicates)±SD. **c,** Effect of cytokine treatment on cell viability of Calu-3 cells treated as in A, as assessed by intracellular ATP levels. Lines represent the mean of n=3±SEM. **d**, SARS-CoV-2 N RNA in the supernatant of SARS-CoV-2 (MOI 0.05, 48h p.i.) infected Calu-3 cells that were left untreated and/or were treated with the indicated amounts of Rapamycin or Bafilomycin A1 (BafA). Lines represent the mean of n=2±SD. **e**, Exemplary standard curve of raw qRT-PCR CT values corresponding to the SARS-CoV-2 RNA copy numbers per ml for one replicate. **f**, Effect of autophagy modulating drugs on cell viability of Calu-3 cells treated as indicated, as assessed by intracellular ATP levels. Lines represent the mean of n=3±SEM. **g**, Exemplary standard curve of raw qRT-PCR CT values corresponding to the SARS-CoV-2 RNA copy numbers per ml for one replicate. **h**, Exemplary CT values of N RNA in the supernatant of SARS-CoV-2 infected samples treated as indicated n=2(technical replicates)±SD for one replicate. Treatment 24 h before infection (Pre), treatments 6 h post infection (Post). See also Figure 5.
